# Carotenogenesis of *Staphylococcus aureus*: new insights and impact on membrane biophysical properties

**DOI:** 10.1101/2020.07.19.210609

**Authors:** Gerson-Dirceu López, Elizabeth Suesca, Gerardo Álvarez-Rivera, Adriana Rosato, Elena Ibáñez, Alejandro Cifuentes, Chad Leidy, Chiara Carazzone

## Abstract

Staphyloxanthin (STX) is a saccharolipid derived from a carotenoid in *Staphylococcus aureus* involved in oxidative-stress tolerance and antimicrobial peptide resistance. In this work, a targeted metabolomics and biophysical study was carried out on native and knock-out *S. aureus* strains to investigate the biosynthetic pathways of STX and related carotenoids. Identification of 34 metabolites at different growth phases (8, 24 and 48h), reveal shifts of carotenoid populations during progression towards stationary phase. Six of the carotenoids in the STX biosynthetic pathway and three menaquinones (Vitamin K_2_) were identified in the same chromatogram. Furthermore, other STX homologues with varying acyl chain structures reported herein for the first time, which reveal the extensive enzymatic activity of CrtO/CrtN. Fourier Transform infrared spectroscopy show that STX increases acyl chain order and shifts the cooperative melting of the membrane indicating a more rigid lipid bilayer. This study shows the diversity of carotenoids in *S. aureus*, and their influence on membrane biophysical properties.

## 1. Introduction

*S. aureus* is a Gram-positive bacterium naturally present in nasal passages and human skin. It is an opportunistic pathogen responsible for nosocomial and acquired infections, including pneumonia, osteomyelitis, meningitis, bacteremia and sepsis (Lowy, 1998; V. Recklinghausen, 2008; Tong et al., 2015). The main concern about this microorganism is the increasing number of resistant strains to different antibiotics (Oldfield and Feng, 2014). Thus, methicillin-resistant *S. aureus* (MRSA) strains pose a serious problems for health, limiting the antimicrobial treatments as reported by the Center for Disease Control and Prevention (CDC), that relates MRSA with more than 323,000 hospitalized patients and 10,600 estimated deaths in the US in 2017 (CDC, 2019). Thereby, an alternative to the use of conventional antibiotics against *S. aureus* has been to employ antimicrobial peptides, such as Daptomycin (Crass et al., 2019; Dhand and Sakoulas, 2014; Heidary et al., 2018; Steenbergen et al., 2005), which act by compromising bacterial membrane integrity. For such molecules, the antimicrobial activity has been shown to depend on how the microorganism modulates the physicochemical properties of its membrane, which include mechanical malleability and lateral diffusion, since they strongly influence the insertion of membrane active peptides. For this reason, antibiotic resistance in *S. aureus* strains has been associated with changes in the membrane composition (Kilelee et al., 2010; Mishra et al., 2009; Xue et al., 2019; Zhang et al., 2014).

The number of studies focused on *S. aureus* membrane lipid composition has increased in recent years. Particularly, a great interest has been directed to a saccharolipid containing carotenoid, known as Staphyloxanthin (STX) (Braungardt and Singh, 2019; Perez-Lopez et al., 2019; Tiwari et al., 2018; Xue et al., 2019), a natural pigment with well-known antioxidant properties (Mishra et al., 2011; Tiwari et al., 2018; Zhang et al., 2018), responsible for the characteristic color of *S. aureus* (Kim and Lee, 2012; Marshall and Wilmoth, 1981a), and associated with tolerance to oxidative stress (Clauditz et al., 2006; Liu et al., 2005; Olivier et al., 2009). However, STX also plays an essential role on the regulation of membrane mechanical properties, and has been shown to hinder the permeability of the membrane to cationic antimicrobial peptides, increasing the virulence and bacterial fitness of *S. aureus* (Crass et al., 2019; Mishra et al., 2011; Vogeser and Zhang, 2018). In the first comprehensive study on STX and other carotenoids of *S. aureus* S41, these pigments were isolated and their chemical structures determined (Marshall and Wilmoth, 1981a), leading to the identification of 17 triterpenic carotenoid compounds and proposal of a biosynthetic pathway (Marshall and Wilmoth, 1981b). However, it was not until 2005 that five of the six enzymes involved in the biosynthesis of STX in *S. aureus* were reported (Pelz et al., 2005). Recently, using *Escherichia coli* mutants (Kim and Lee, 2012), a sixth enzyme involved in the biosynthetic pathway of STX was reported (**Fig. 1a**). In these studies, open column chromatography (OCC) or thin-layer chromatography (TLC) were mostly used as the separation step prior to the mass spectrometric (MS) analysis of carotenoids. In addition to these carotenoids, other secondary metabolites exhibit structural similarities to STX related biproducts. These include menaquinones (MK) or vitamin K_2_, unsaturated polyisoprenes of 2-methyl-1,4-naphthoquinones, involved in the aerobic or anaerobic respiration of *S. aureus*, thanks to the possible transfer of two electrons (Kurosu and Begari, 2010; Wakeman et al., 2012). Menaquinones are more non-polar compounds due to the lack of conjugated double bonds. *S. aureus* shows three types of menaquinones with different length of the aliphatic chain (**Fig. 2a**), MK (n=7, 8 and 9) (Marshall and Wilmoth, 1981a; Taylor’ and Bavies, 1983; Wakeman et al., 2012). In this study, we have included the identification of these compounds to provide a more complete picture of carotenoid related metabolites present in the *S. aureus* membrane.

**Figure 1.**
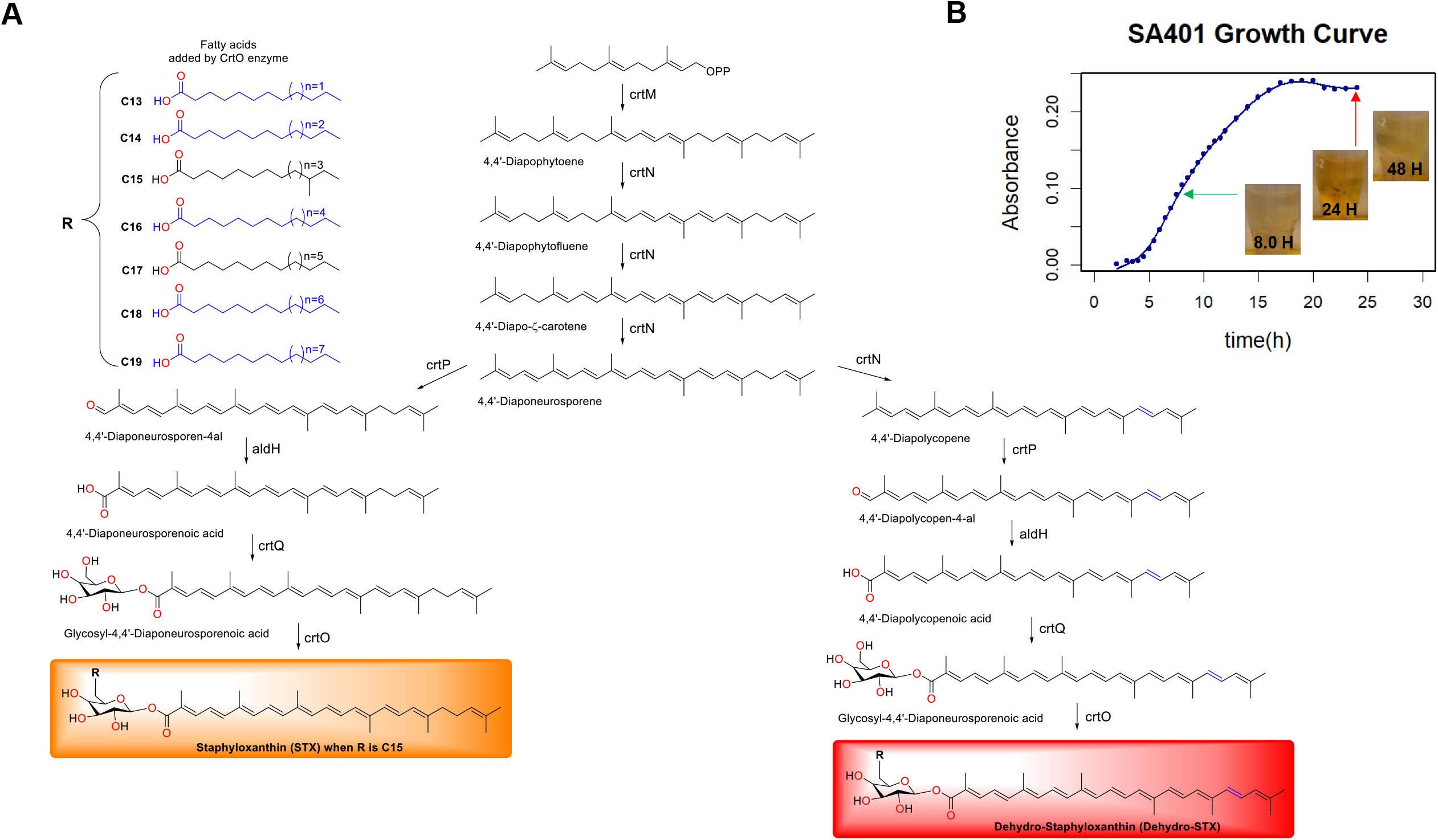
STX biosynthetic pathway and growth curve of S. aureus. (A) Main (right) and alternative (left) pathways including the variation of fatty acids, adapted from (Kim and Lee, 2012). (B) Growth curve for SA401 *S. aureus* strain for determinate the exponential and stationary phases.

**Figure 2.**
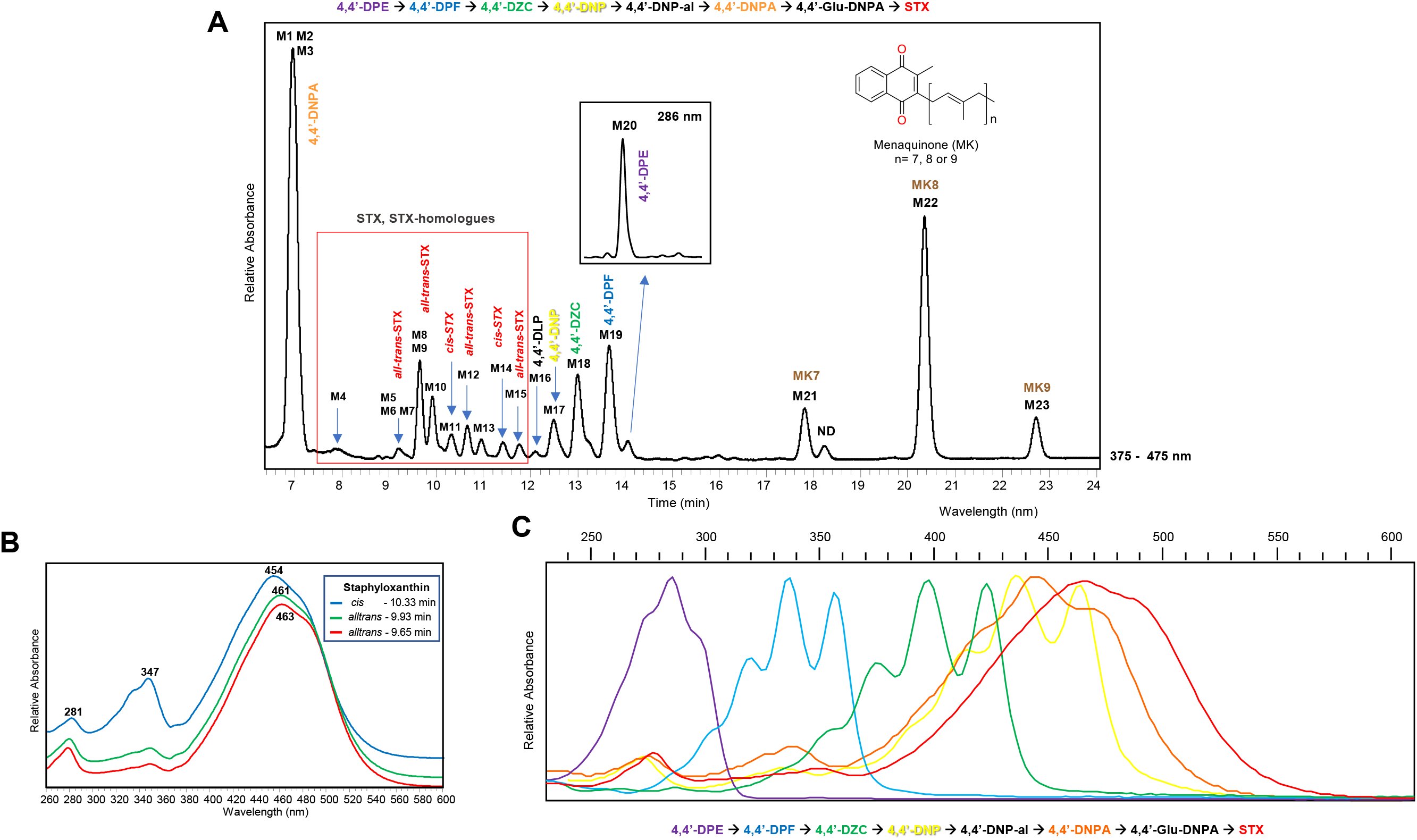
HPLC-DAD analysis. (A) Chromatogram extracted in the UV-visible region (375-475 nm and 286 nm). (B). Absorbances of *cis* and *trans* STX. (C) Bathochromic shift in the absorbances of six carotenoids involved in STX biosynthetic pathway.

Previously, the fatty acids composition and headgroup composition of *S. aureus* phospholipids have been described as a mechanism for modulating the biophysical properties of the bacterial membrane, showing an influence on the pathogenicity and resistance to antimicrobial peptides (Hernández-Villa et al., 2018; Kilelee et al., 2010; Mishra and Bayer, 2013; Sen et al., 2016; Zhang et al., 2014). Hence, biophysical studies on membrane stiffness in *S. aureus* have been developed by fluorescence spectroscopy using the probe 1,6-diphenyl-1,3,5-hexatriene (DPH) (Mishra et al., 2011; Perez-Lopez et al., 2019; Sen et al., 2016; Tiwari et al., 2018). However, the insertion of fluorescent probes into the membrane is cumbersome, with a high risk of incorporation into the cell wall, therefore affecting the proper report of the membrane behavior. Instead, Fourier-transform infrared spectroscopy (FT-IR) provides a direct approach for studying the biophysical behavior *in vivo* of the lipids present in bacterial membranes (Ocampo et al., 2010; Schultz and Naumann, 1991). The CH_2_ stretch vibration of phospholipids is a dominant signal in bacterial cells and reflects directly the physical properties of the lipids that compose the bilayer (Scherber et al., 2009). The thermotropic shift of the CH_2_ stretch indicates cooperative changes in the lipid packing behavior of the membrane as the membrane moves from a tightly packed gel-like phase (L_β_) at low temperatures to a more mobile liquid-crystalline phase at high temperatures close to its growth temperature (Ocampo et al., 2010). *S. aureus* presents a reproducible cooperative shift in the CH_2_ wavenumber around 15°C, suggesting that below 15°C the membrane of *S. aureus* has a significantly higher rigidity. Shifts in the wavenumber and this cooperative event are therefore used as indications of changes in the rigidity of the bilayer membrane (Ocampo et al., 2010).

In the present study, we conduct an in-depth characterization of the carotenoids of *S. aureus* at different stages of cell growth, in order to better understand their biosynthetic routes and their impact on Staphyloxanthin production and on membrane biophysical properties. For the identification of carotenoids and structurally related compounds, a robust methodology based on HPLC-DAD-MS/MS analysis was proposed. Complementary structural information was obtained from UV-visible absorbances and tandem-mass spectrometry-based detection systems. Further metabolites coverage was obtained applying both APCI and ESI ionization sources. We present the versatility of a C18 column in the characterization of the carotenoids and STX-homologues compounds in wild type and mutant strains of *S. aureus*, complemented by the usefulness of a C30 column for the characterization of new Dehydro-STX-FA, through a simple and reliable method, without the need for previous TLC or OCC. In addition, we included analysis of the shift in the CH_2_ stretch wave number by FTIR to determine the impact of these carotenoids on the biophysical properties of the bacterial membrane without the need of exogenous fluorescent probes.

## 2. Experimental

### 2.1 Chemicals and reagents

HPLC-grade methanol, acetonitrile, ethyl acetate, and methyl tert-butyl ether (MTBE) were purchased from Honeywell (Michigan, USA), J.T. Baker (Palo Alto, CA, USA) and VWR (Leuven, Belgium), respectively. HPLC-grade acetic acid and ammonium acetate were purchased from Fluka (St. Louis, MO, USA). Butylated hydroxytoluene (BHT) and NaCl ReagentPlus (>99%) were purchased from Sigma-Aldrich (St. Louis, MO, USA). Tryptone (OXOIO, Basigstoke, Hampshire, England), NaCl (ACS, J.T. Baker, USA), and a yeast extract (Dibico, México D.F., México) were used for LB medium preparation. HPLC-grade water was obtained from a water purification system Heal Force Smart-Mini (Shangai, China) and Milli-Q system (Millipore, Billerica, MA, USA).

### 2.2 Bacterial growth conditions

Two clinical methicillin-susceptible *Staphylococcus aureus* strains were used. The strain denominated SA401 was provided by CIMIC (Centro de Investigaciones Microbiologicas of Universidad de los Andes, Bogotá D.C., Colombia) and a full description of its biophysical properties have been published (Perez-Lopez et al., 2019), whereas SA144 strain was obtained from the Center for Molecular and Translational Human Infectious Diseases Research (Houston, USA). In addition, a crtM deletion mutant of SA144 (SA145) and a crtMN plasmid complement variant of SA145 (SA147) that recovered the carotenoid biosynthesis, were used (Mishra et al., 2011). Also, *S. aureus* subsp. aureus Rosenbach ATCC® 25923™ strain was analyzed as well. One colony of each *S. aureus* strain was grown overnight (37°C), under constant agitation (250 rpm), in 10 mL of LB medium containing (per liter) 10 g of Tryptone, 10 g of NaCl, and 5 g of the yeast extract. SA147 was grown on LB medium containing erythromycin 1ug/ml. Then, the cells were diluted (1:1000) in flasks containing 150 ml of fresh LB medium and cultivated for 8, 24 or 48 hours. Subsequently, the cells were harvested by centrifuging for 10 min at 8500 rpm at 4 °C (Thermo Scientific, USA), and the pellet was frozen (−80°C) and lyophilized for almost 24 hours (LABCONCO, Kansas City, MO, USA).

### 2.3 Cellular growth curves

After growing, optical densities (OD) measures were obtained from the main culture of *S. aureus* strain SA401, which was inoculated with 1:1000 of overnight culture and incubated at 37°C until full stationary phase (24 h) was reached (A600=0.20-0.25). Optical densities were measured in triplicates at 600nm with a NanoDrop 2000 UV-Vis spectrophotometer (Thermo Scientific, Wilmington, DE, USA), every 30 minutes for the first 12 hours and every hour during the next 12 hours. Once the full stationary phase was reached, the 48 hours of growth was considered the late stationary phase.

### 2.4 Carotenoids extraction

The extraction of carotenoids was achieved using a modified version of the Marshall method (Marshall and Wilmoth, 1981a). Briefly, 100 mg of lyophilized cells were accurately weighed in a falcon tube containing 10 glass beads, dissolved in 2.0 mL of methanol containing BHT (0.1%, w/v) and vortex-mixed for 5 min. After centrifugation at 8500 rpm for 10 min, the supernatant containing the pigments was gently aspirated with a glass Pasteur pipette and the extraction was repeated twice with 1.0 mL of MeOH each time. All methanolic phases containing the carotenoids were combined, successively shaken with ethyl acetate and 1.7M NaCl (1:3 v/v), and centrifuged again at 8500 rpm for 5 min. Successively, the upper organic phase was carefully drawn, dried with anhydrous Na_2_SO_4_, decanted into an amber glass tube, and finally dried with nitrogen gas. The extracts are removed from the triglycerides (TAGs) that affect the ionization of carotenoids in LC-MS, using low temperatures (−20) (Mariutti and Mercadante, 2018; Marshall and Wilmoth, 1981a).

### 2.5 HPLC-DAD-APCI-MS/MS analysis

The extracts obtained were resuspended and analyzed by high-performance liquid chromatography with diode array detection and atmospheric pressure chemical ionization tandem mass (HPLC-DAD-APCI-MS/MS), using an Agilent 1100 series liquid chromatograph equipped with a binary pump, online degasser, and autosampler (Santa Clara, CA, USA) coupled to an Ion Trap Mass Spectrometer through APCI operated in positive and negative ionization mode (Agilent ion trap 6320, Agilent Technologies, Santa Clara, CA, USA). The instrument was controlled by LC ChemStation 3D Software Rev. B.04.03 (Agilent Technologies, Santa Clara, CA, USA). All extracts were dissolved (10-40 mg mL^−1^) in pure MeOH or ACN (0.1% acetic acid) and filtered using 0.45 μm nylon filters prior to analysis. The RP-HPLC separation was carried out at room temperature with 10 to 20 µL injection volume on a Zorbax SB-C18 column (150 mm × 4.6 mm i.d., 3.5 µm particle size, Agilent Technologies, Santa Clara, CA, USA), and a YMC-C30 reversed-phase column (150 × 4.6 mm i.d., 3 μm particle size; YMC Europe, Schermbeck, Germany). A precolumn YMC-C30 (10 × 4 mm, 5 μm particle size) was used for the analysis. The mobile phases used were ammonium acetate 400 mg/L in a solvent mixture of methanol: methyl tert-butyl ether: water (80:18:2 v/v/v for solution A and 8:90:2 v/v/v for solution B). The elution gradient in C18 column at a constant flow rate of 300 µL/min was as follows: 5% B in the first 3 minutes, after from 5% to 13% B in 9 minutes, from 13% to 100% B in 7 minutes and an isocratic hold at 100% B for 4 minutes. Final reconditioning from 100% to 5% solution B in 2 minutes and then maintained isocratically for 9 minutes (Hrvolová et al., 2016; Schex et al., 2018). For C30 column the elution gradient was: 5% B in the first 3 minutes, after from 5% to 13% B in 9 minutes, from 13% to 25% B in 7 minutes, from 25% to 100% B in 4 minutes and an isocratic hold at 100% B for 2 minutes. Final reconditioning from 100% to 5% solution B in 2 minutes and then maintained isocratically for 9 minutes at a constant flow rate of 500 µL/min. The DAD recorded at 230, 330, 460, and 490 nm, although spectra from 190 to 700 nm were also obtained (peak width 0.1 min (2 s), slit 4 nm).

The APCI source was operated with the following parameters: drying temperature, 350 °C; vaporizer temperature, 400 °C; drying gas flow rate, 7 L/min; capillary voltage, −3.6 kV; nebulizer gas pressure, 45 psi; corona current, 4000 nA. Full scan spectra were obtained in the range from m/z 150 to 1200 (Hrvolová et al., 2016; Novotny et al., 2005). Untargeted and targeted MS/MS data dependent-scans were carried out, fragmenting the two highest precursor ions (10000 counts threshold; 1 V Fragmentor amplitude).

### 2.6 HPLC-DAD-ESI-MS/MS analysis

In order to widen the range of detectable carotenoids and menaquinones observed with the APCI source, the extracts were also analyzed by HPLC-DAD-MS/MS with electrospray ionization (ESI) using a ultra-high-performance liquid chromatographer Dionex UltiMate 3000 equipped with a binary pump, online degasser, autosampler, and a thermostated column compartment coupled with an LCQ Fleet™ Ion Trap Mass Spectrometer through ESI source operated in positive mode (Thermo Scientific, San Jose, CA, USA). Raw metabolite data were acquired and processed using the Xcalibur 3.0 software (Thermo Scientific, San Jose, CA, USA). The RP-HPLC separation was carried out at 30°C with 10 µL injection volume, using the same gradients described in section 2.5. Diode-array detection was performed over the entire UV-vis range (190 - 800 nm), and the characteristic absorbances of the carotenoids were extracted between 230-550 nm.

MS operating conditions were previously optimized using flow injection analysis of a 10-ppm solution of the carotenoid extract in ACN (0.1% acetic acid). The ESI source was operated with the following parameters: ionization voltage 5.5 kV, capillary temperature of 330°C, sheath gas flow rate of 9 arbitrary units, auxiliary gas flow rate of 2 arbitrary units. The ion trap was set to operate in full scan (*m/z* 65-1200 mass range), and data-dependent MS/MS (30% collision energy) mode to obtain the corresponding fragment ions with an isolation width of 3 m/z.

### 2.7 FTIR measurements

FTIR analysis was performed according to a previously described method (Ocampo et al., 2010), with some modifications. Briefly*, S. aureus* strains were grown in the same conditions described in section 2.2. Cells were measured from inoculations (1:1000) overnight culture and incubated for 18 hours. Subsequently, cells were washed with 30 mL phosphate-buffered saline (PBS) solution and centrifugated. Cell pellet was smeared onto Ge windows and placed within an adapted-built Peltier temperature controller inside of the FTIR chamber (IRTracer-100, Shimadzu, Japan). Temperature was ramped between 5 °C and 50°C, performing a scan from 4000 to 400 cm^−1^, with a resolution of 1 cm^−1^ and 80 spectrograms for each temperature point. Analysis was carried out for data between 2860 and 2840 cm^−1^ where the CH_2_ symmetric stretch vibration is centered, enabling to characterize the thermotropic chain melting behavior of native bacterial membranes from *S. aureus*. For every temperature, data was fitting to a polynomial function using R (RStudio, 2020). With the same program, pick position in the spectrograms and derivates were determined.

## 3. Results

### 3.1. Establishing S. aureus growth phases and carotenoids extraction

In order to monitor the progression of carotenoid species in *S. aureus* during STX biosynthesis at different growth stages, a bacterial growth curve of SA401 strain (wild type) was first measured to determine the exponential (8 hours), full stationary (24 hours) and late stationary (48 hours) growth stages, as depicted in Fig. 1b. This information is useful to understand the metabolic change in the different growth stages, observed by comparing chromatographic profiles of these phases, as discussed in the end of Section 3.2.

For the extraction of carotenoids, several factors such as light exposure and temperature were carefully controlled to avoid degradation. Reported methods frequently involve the use of different solvents, such as methanol, acetone, and ethyl acetate (Kim and Lee, 2012; Marshall and Wilmoth, 1981a). In this work, a modified version of the Marshall method was used, changing incubation with MeOH heated in a water bath at 55°C (Marshall and Wilmoth, 1981a) by a maceration step using vortexing with glass beads for 5 minutes (Hewelt-Belka et al., 2014), to improve the cell lysis (Hartz et al., 2018; Ye et al., 2006), and prevent thermal degradation of carotenoids. The resulting crude extracts showed an orange color of high intensity (24 hours), indicating the presence of carotenoid derivatives. Carotenoid extracts from SA401 strain were obtained at different growth stages for 8, 24, and 48 hours (**Fig. 1b**), whereas ATCC, SA144, SA145 and SA147 strains were all grown for 24 hours before extraction. Experiments were carried out in triplicate, following the same extraction procedure.

### 3.2. Carotenoids profiling by HPLC-DAD-(APCI/ ESI)-MS/MS analysis

Different stationary phases, buffer solutions and gradients were tested to optimize chromatographic resolution before HPLC-DAD-MS/MS analysis (Chu et al., 2011; Kim and Lee, 2012; Mijts et al., 2005; Pelz et al., 2005). Although C30 columns are commonly used for carotenoids analysis due to the higher resolution capacity for these large molecular size terpenoids, improved chromatographic separation for a larger number of target metabolites involved in STX biosynthesis were found with a C18 column, employing a typical mobile phase for carotenoids analysis based on MTBE:MeOH:Water with ammonium acetate (Amorim-Carrilho et al., 2014; Hrvolová et al., 2016; Novotny et al., 2005; Schex et al., 2018), as described in the experimental section. Nevertheless, both C18 and C30 stationary phases showed complementary information in the metabolomic analysis. Selecting specific wave lengths for xanthophylls, carotenes, and menaquinones in the UV-visible region (230 - 520 nm), a total of 34 metabolites were detected in the extracts of *S. aureus* after 8 h and 24 h of growth (**Fig. 2a**). The identity of the detected metabolites could be confidently assigned by comparing UV-visible absorbances and MS/MS spectral information obtained using both APCI and ESI ionization sources. Table 1 summarizes the tentatively identified metabolites (M1-M34), including their retention time, ionization mode as well as MS and UV-visible spectral information. The analyzed *S. aureus* extracts allowed the identification of a large family of carotenoids attached to a saccharolipid residue, including STX (M3-M5, M8, M10, and M11), STX-homologues (M6, M7, M9, M12-M15, M24, M27, and M34), as well as Dehydro-STX (M26, M28 and M32) and Dehydro-STX-homologues (M29, M30, M31, and M33). In addition, biosynthetic precursors, such as hydrocarbon carotenes (M16-M20) and carotenoid acids (M1 and M2), as well as the three terpenoid derivatives such as menaquinones (M21-M23) were also characterized (see Table 1).

**Table 1.** Carotenoids and menaquinones identified in *S. aureus*.

| Metabolite number | Ret. time (min) | Identified compound | Abbreviation | Molecular formula | Ionization | Molecular ion (m/z) | MS/MS product ions (m/z) in ESI | MS/MS product ions (m/z) in APCI | UV-Vis max wavelength (nm) |
| --- | --- | --- | --- | --- | --- | --- | --- | --- | --- |
| M1 | 6.65 | 4,4'-diapolycopenoic acid | 4,4'-DLPA | C<sub>30</sub>H<sub>38</sub>O<sub>2</sub> | APCI (-) | [M+H]<sup>+</sup> / 429.4 [M-H]<sup>-</sup> | ND | 385, 279, 227 | 446, 474, 500 |
| M2 | 6.86 | 4,4'-diaponeurosporenoic acid | 4,4'-DNPA | C<sub>30</sub>H<sub>40</sub>O<sub>2</sub> | ESI (+) / APCI (-) | 433.3 [M+H]<sup>+</sup> / 431.7 [M-H]<sup>-</sup> | 415, 387, 363, 340, 309, 288, 274, 267. | 387, 267, 229 | (420), 447, 472 |
| M3 | 7.00 | Staphyloxanthin-Isomer1 | STX-Iso1 | C<sub>51</sub>H<sub>78</sub>O<sub>8</sub> | APCI (-) | 877.7 [M+AcOH-H]<sup>-</sup> | ND | 818 (M<sup>-</sup>), 758, 473, 431, 387, 255. | 465 (488) |
| M4 | 7.60 | Staphyloxanthin-Isomer2 | STX-Iso2 | C<sub>51</sub>H<sub>78</sub>O<sub>8</sub> | APCI (-) | 877.7 [M+AcOH-H]<sup>-</sup> | ND | 818 (M<sup>-</sup>), 758, 473, 431, 387, 255. | 465 (488) |
| M5 | 8.80 | Staphyloxanthin-Isomer3 | STX-Iso3 | C<sub>51</sub>H<sub>78</sub>O<sub>8</sub> | APCI (-) | 877.7 [M+AcOH-H]<sup>-</sup> | ND | 818 (M<sup>-</sup>), 758, 473, 431, 387, 255. | 465 (488) |
| M6 | 8.80 | Tridecanoyl-glucosyl-4,4'-4,4'-diaponeurosporenoic acid-Isomer1 | STX-C13-Iso1 | C<sub>49</sub>H<sub>74</sub>O<sub>8</sub> | APCI (-) | 849.7 [M+AcOH-H]<sup>-</sup> | ND | 789,6 (M-H<sup>-</sup>); 531, 471, 431, 387. | 463 (488) |
| M7 | 9.15 | Tridecanoyl-glucosyl-4,4'-4,4'-diaponeurosporenoic acid-Isomer1 | STX-C13-Iso2 | C<sub>49</sub>H<sub>74</sub>O<sub>8</sub> | APCI (-) | 849.7 [M+AcOH-H]<sup>-</sup> | ND | 789,6 (M-H<sup>-</sup>); 531, 471, 431, 387. | 463 (488) |
| M8 | 9.65 | Staphyloxanthin-Isomer4 | STX-Iso4 | C<sub>51</sub>H<sub>78</sub>O<sub>8</sub> | ESI (+) / APCI (-) | 818.3 [M]<sup>+</sup>; 841.4 [M+Na]<sup>+</sup> / 877.7 [M+AcOH-H]<sup>-</sup> | M<sup>2</sup>[818.3] => 800,749, 726, 432, 415, 387, 340. M<sup>2</sup>[841.4] => 823, 735, 456, 410. | 818 (M<sup>-</sup>), 758, 473, 431, 387, 255. | 463, (488) |
| M9 | 9.65 | Heptadecanoyl-glucosyl-4,4'-diaponeurosporenoic acid-Isomer1 | STX-C17-Iso1 | C<sub>53</sub>H<sub>82</sub>O<sub>8</sub> | APCI (-) | 905.6 [M+AcOH-H]<sup>-</sup> | ND | 845,4 (M-H<sup>-</sup>); 431; 387; 269,5. | 463 (488) |
| M10 | 9.93 | Staphyloxanthin-Isomer5 | STX-Iso5 | C<sub>51</sub>H<sub>78</sub>O<sub>8</sub> | ESI (+) / APCI (-) | 818.3 [M]<sup>+</sup>; 841.4 [M+Na]<sup>+</sup> / 877.7 [M+AcOH-H]<sup>-</sup> | M<sup>2</sup>[818.3] => 800,749, 726, 432, 415, 387, 340. M<sup>2</sup>[841.4] => 823, 735, 456, 410. | 818 (M<sup>-</sup>), 758, 473, 431, 387, 255. | 463, (488) |
| M11 | 10.33 | Staphyloxanthin-Isomer6 | STX-Iso6 | C<sub>51</sub>H<sub>78</sub>O<sub>8</sub> | ESI (+) / APCI (-) | 818.3 [M]<sup>+</sup>; 841.4 [M+Na]<sup>+</sup> / 877.7 [M+AcOH-H]<sup>-</sup> | M<sup>2</sup>[818.3] => 800,749, 726, 432, 415, 387, 340. M<sup>2</sup>[841.4] => 823, 735, 456, 410. | 818 (M<sup>-</sup>), 758, 473, 431, 387, 255. | 463, (488) |
| M12 | 10.66 | Heptadecanoyl-glucosyl-4,4'-diaponeurosporenoic acid-Isomer2 | STX-C17-Iso2 | C<sub>53</sub>H<sub>82</sub>O<sub>8</sub> | APCI (-) | 905.6 [M+AcOH-H]<sup>-</sup> | ND | 831,4 (M-H<sup>-</sup>); 431; 387; 311; 255. | 463 (488) |
| M13 | 10.97 | Heptadecanoyl-glucosyl-4,4'-diaponeurosporenoic acid-Isomer3 | STX-C17-Iso3 | C<sub>53</sub>H<sub>82</sub>O<sub>8</sub> | APCI (-) | 905.6 [M+AcOH-H]<sup>-</sup> | ND | 831,4 (M-H<sup>-</sup>); 431; 387; 311; 255. | 463 (488) |
| M14 | 11.76 | Octadecanoyl-glucosyl-4,4'-diaponeurosporenoic acid | STX-C18 | C<sub>54</sub>H<sub>84</sub>O<sub>8</sub> | APCI (-) | 919.8 [M+AcOH-H]<sup>-</sup> | ND | 859,6 (M-H<sup>-</sup>); 431; 387; 283. | 463 (488) |
| M15 | 11.41 | Nonadecanoyl-glucosyl-4,4'-diaponeurosporenoic acid | STX-C19 | C<sub>55</sub>H<sub>86</sub>O<sub>8</sub> | APCI (-) | 933.9 [M+AcOH-H]<sup>-</sup> | ND | 873,5 (M-H<sup>-</sup>); 431; 387; 297. | 463 (488) |
| M16 | 12.11 | 4,4'-Diapolycopene | 4,4'-DLP | C<sub>30</sub>H<sub>40</sub> | ESI (+) / APCI (+) | 400.3 [M+H]<sup>+</sup> / 401.5 [M+H]<sup>+</sup> | 385, 357, 308 | ND | 441, 465, 496 |
| M17 | 12.29 | 4,4'-Diaponeurosporene | 4,4'-DNP | C<sub>30</sub>H<sub>42</sub> | ESI (+) / APCI (+) | 402.4 [M+H]<sup>+</sup> / 403.5 [M+H]<sup>+</sup> | 388, 334, 310, 242. | 386, 356, 346, 327, 291, 267, 187, 148. | 412; 435; 464 |
| M18 | 13.00 | 4,4'-Diapo-ζ-carotene | 4,4'-DZC | C<sub>30</sub>H<sub>44</sub> | APCI (+) | 405.5 [M+H]<sup>+</sup> | ND | 386, 362, 336, 295, 225, 173, 157. | 382; 399; 423 |
| M19 | 13.66 | 4,4'-Diapophytofluene | 4,4'-DPF | C<sub>30</sub>H<sub>46</sub> | APCI (+) | 407.5 [M+H]<sup>+</sup> | ND | 388, 349, 336, 323, 295, 225, 173, 159. | 330; 347; 366 |
| M20 | 14.07 | 4,4'-Diapophytoene | 4,4'-DPE | C<sub>30</sub>H<sub>48</sub> | APCI (+) | 409.5 [M+H]<sup>+</sup> | ND | 392, 368, 354, 326, 299, 286, 215, 159. | 275; 286; 297 |
| M21 | 17.82 | Menaquinone 7 | MK7 | C<sub>46</sub>H<sub>64</sub>O<sub>2</sub> | ESI (+) / APCI (+) | 649.4 [M+H]<sup>+</sup> / 649.7 [M+H]<sup>+</sup> | 592, 567, 442, 291, 265, 225, 187. | 632, 594, 568, 464, 227, 187. | 247, 269, 329 |
| M22 | 20.36 | Menaquinone 8 | MK8 | C<sub>51</sub>H<sub>72</sub>O<sub>2</sub> | ESI (+) / APCI (+) | 717.4 [M+H]<sup>+</sup> / 717.7 [M+H]<sup>+</sup> | 699, 635, 593, 531, 291, 265, 227, 225. | 700, 649, 636, 532, 500, 403, 227. | 247, 269, 329 |
| M23 | 22.72 | Menaquinone 9 | MK9 | C<sub>56</sub>H<sub>80</sub>O<sub>2</sub> | ESI (+) / APCI (+) | 785.4 [M+H]<sup>+</sup> / 785.8 [M+H]<sup>+</sup> | 767, 717, 649, 633, 599, 581, 525, 291, 265, 225. | 744, 730, 718, 704, 636, 600, 568, 500, 227. | 247, 269, 329 |
| M24* | 14.76* | Tetradecanoyl-glucosyl-4,4'-diaponeurosporenoic acid | STX-C14 | C<sub>50</sub>H<sub>76</sub>O<sub>8</sub> | APCI (-) | 863.6 [M+AcOH-H]<sup>-</sup> | ND | 803.4 (M-H)<sup>-</sup>, 531, 471, 431, 387. | 430, 460, 483 |
| M25* | 15.30* | Heptadecanoyl-glucosyl-4,4'-diaponeurosporenoic acid-Isomer4 | STX-C17-Iso4 | C<sub>53</sub>H<sub>82</sub>O<sub>8</sub> | APCI (-) | 905.6 [M+AcOH-H]<sup>-</sup> | ND | 845.5 (M-H)<sup>-</sup>, 603. | 466 (488) |
| M26* | 17.35* | Pentadecanoyl-glucosyl-4,4'-diapolycopenoic acid-Isomer1 | Dehydro-STX-C15-Iso1 | C<sub>51</sub>H<sub>76</sub>O<sub>8</sub> | APCI (-) | 875.7 [M+AcOH-H]<sup>-</sup> | ND | 815.6 (M-H)<sup>-</sup>; 429; 385; 283; 255; 241. | 455, 483, 512 |
| M27* | 17.70* | Hexadecanoyl-glucosyl-4,4'-diaponeurosporenoic acid | STX-C16 | C<sub>52</sub>H<sub>80</sub>O<sub>8</sub> | APCI (-) | 891.7 [M+AcOH-H]<sup>-</sup> | ND | 831.6 (M-H)<sup>-</sup>; 431; 387; 255.3. | 430, 460 (488) |
| M28* | 18.57* | Pentadecanoyl-glucosyl-4,4'-diapolycopenoic acid-Isomer2 | Dehydro-STX-C15-Iso2 | C<sub>51</sub>H<sub>76</sub>O<sub>8</sub> | APCI (-) | 875.7 [M+AcOH-H]<sup>-</sup> | ND | 815.6 (M-H)<sup>-</sup>; 429; 385; 283; 255; 241. | 455, 483, 512 |
| M29* | 19.37* | Heptadecanoyl-glucosyl-4,4'-diapolycopenoic acid-Isomer1 | Dehydro-STX-C17-Iso1 | C<sub>53</sub>H<sub>80</sub>O<sub>8</sub> | APCI (-) | 903.6 [M+AcOH-H]<sup>-</sup> | ND | 843.6 (M-H)<sup>-</sup>; 429; 387.5; 283, 269. | 455, 483, 512 |
| M30* | 19.44* | Tridecanoyl-glucosyl-4,4'-diapolycopenoic acid | Dehydro-STX-C13 | C<sub>49</sub>H<sub>72</sub>O<sub>8</sub> | APCI (-) | 847.6 [M+AcOH-H]<sup>-</sup> | ND | 787.6 (M-H)<sup>-</sup>; 531, 471, 431, 387. | 455, 483, 512 |
| M31* | 19.60* | Heptadecanoyl-glucosyl-4,4'-diapolycopenoic acid-Isomer2 | Dehydro-STX-C17-Iso2 | C<sub>53</sub>H<sub>80</sub>O<sub>8</sub> | APCI (-) | 903.6 [M+AcOH-H]<sup>-</sup> | ND | 843.6 (M-H)<sup>-</sup>; 429; 387.5; 283, 269. | 455, 483, 512 |
| M32* | 19.86* | Pentadecanoyl-glucosyl-4,4'-diapolycopenoic acid-Isomer3 | Dehydro-STX-C15-Iso3 | C<sub>51</sub>H<sub>76</sub>O<sub>8</sub> | APCI (-) | 875.7 [M+AcOH-H]<sup>-</sup> | ND | 815.6 (M-H)<sup>-</sup>; 429; 385; 283; 255; 241. | 455, 483, 512 |
| M33* | 20.55* | Heptadecanoyl-glucosyl-4,4'-diapolycopenoic acid-Isomer3 | Dehydro-STX-C17-Iso3 | C<sub>53</sub>H<sub>80</sub>O<sub>8</sub> | APCI (-) | 903.8 [M+AcOH-H]<sup>-</sup> | ND | 843.6 (M-H)<sup>-</sup>; 429; 387.5; 283, 269. | 455, 483, 512 |
| M34** | 21.15** | Eicosanoyl-glucosyl-4,4'-diaponeurosporenoic acid | STX-C20 | C<sub>56</sub>H<sub>88</sub>O<sub>8</sub> | APCI (-) | 947.7 [M+AcOH-H]<sup>-</sup> | ND | 887.6 (M-H)<sup>-</sup>; 431; 387; 311. | ND |

\*Compounds present only Strain SA147 and retention time in C30 column Figure 4b
\*\*Compound observed in Strain SA401, retention time in C30 column.

The MS fragmentation pattern of carotenoid acids, such as 4,4’-diaponeurosporenoic acid (4,4’-DNPA) and 4,4’-diapolycopenoic acid (4,4’-DLPA), is mainly characterized by the loss of HCOOH (−46 amu) or CO_2_ (−44 amu), in positive and negative ionization mode, respectively. STX is a glycosylated 4,4’-DNPA bounded to a C15 fatty acid (FA), which can be represented as FA-Glu-4,4’-DNPA. The MS/MS fragmentation of STX and other identified biosynthetic homologues is characterized by the loss of the fatty acid-glucose (FA-Glu) residue. Thus, STX and STX-homologues show the characteristic *m/z* 431 [M-FA-Glu-H]^−^ product ions in negative ionization mode, whereas Dehydro-STX and Dehydro-STX homologues exhibit *m/z* 429 as major product ion. The different fatty acids (FA) attached to the STX core (Glu-4,4’-DNPA) or Dehydro-STX core (Glu-4,4’-DLPA) could also be confirmed by minor product ions in the MS/MS spectra. Free FA were also found in the first minutes of the chromatographic separation (3-4.5 min) at C18 column.

LC-MS^n^ analysis of carotenoid extracts with an APCI source allowed the identification of the thirty metabolites involved in the carotenoid biosynthetic pathway in *S. aureus*. The carotenoid acids 4,4’-DLPA (M1) at *m/z* 429 [C_30_H_38_O_2_-H]^−^ (in SA147) (Kim and Lee, 2012) and 4,4’-DNPA (M2) at *m/z* 431 [C_3_0H_40_O_2_-H]^−^ presented characteristic loss of CO_2_ in APCI(−) mode. The metabolite 4,4’-DNPA was also detected at *m/z* 433 [C_30_H_40_O_2_+H]^+^ in APCI(+), showing MS/MS spectrum peaks at *m/z* 415 [M+H−18]^+^ and *m/z* 387 [M+H-46]^+^, which indicate the loss of H_2_O and HCOOH, respectively (Figure S1) (Kim and Lee, 2012; Marshall and Wilmoth, 1981a; Pelz et al., 2005). Similar ions were observed in ESI (+). In addition, this carotenoid acid presented absorbances at (420 nm), 446 nm, and 472 nm, similar to those reported in methanol (Marshall and Wilmoth, 1981a).

In the case of STX and STX-homologues, two major ions were observed in the mass spectrum in APCI (−). STX isomers (M3, M4, M5, M8, M10 and M11) exhibited *m/z* 817.8 [M-H]^−^ (Kim and Lee, 2012) and *m/z* 877.8 [M+AcOH-H]-, as deprotonated molecular ions and the STX adduct with acetic acid (**Fig. 3a**), respectively. Acetic acid from the mobile phase is presumably attached to the hydroxyl groups of the glucose (Amorim-Carrilho et al., 2014; Hrvolová et al., 2016; Novotny et al., 2005; Schex et al., 2018). Different lipid chains, ranging from C13 to C20, attached to the STX core (Glu-4,4’-DNPA) evidenced the broad variety of the STX homologues in this biosynthetic pathway. Thus, STX-homologues such as STX-C13 (M6, M7) at *m/z* 849.7, STX-C14 (M24) at *m/z* 863.7, STX-C16 (M27) at *m/z* 891.7, STX-C17 (M9, M12, M13, M25) at *m/z* 905.6, STX-C18 (M14) at *m/z* 919.7, STX-C19 (M15) at *m/z* 933.8, and STX-C20 (M34) at *m/z* 947.6 were assigned by the [M+AcOH-H]^−^ adduct ion, see table 1 and figure S2. Besides, MS/MS analysis showed characteristic *m/z* 431 and *m/z* 387, corresponding to the loss the fatty acid-glucose (FA-Glu) residue, which demonstrate the structural similarity of STX-homologues with 4,4’-DNPA. In addition, free FAs observed in the first minutes of the chromatogram confirm the wide diversity of these molecules bonded to the STX core, as previously stated (**Fig. 3b**). STX was found in ESI(+) as the ions previously reported in the literature (Marshall and Wilmoth, 1981a; Pelz et al., 2005). Thus, the molecular ion at *m/z* 818.3 [C_51_H_78_O_8_]^+^ and the sodium adduct at *m/z* 840.4 [C_51_H_78_O_8_+Na]^+^, addition typical fragments associated to STX were observed (Table 1). Both 4,4’-DNPA and STX lose a toluene molecule (M+H-92) and (M-386-92) respectively, generating a fragment at *m/z* 340, characteristic of carotenoids (Amorim-Carrilho et al., 2014). Additionally, the *m/z* 749 [M-69]^+^ ion was generated by the loss of an isopentenyl fragment (Figure S3). Similarly, these ions were observed in MS/MS experiments performed on *m/z* 819 [M+H]^+^ by flow injection analysis of the crude carotenoid extract (data not shown). Furthermore, different types of visible absorbances were observed for STX and STX-homologues peaks, detected between 9.21 and 11.76 minutes, which allowed the classification as *cis/trans* isomers (**Fig. 2a**). Thus, the peaks at 10.33 and 11.41 minutes presented absorption in visible region (453-454 nm) and an additional peak in the UV region (347 nm), typically called *cis* peak (**Fig. 2b**) (O’neil and Schwartz, 1992), whereas the other peaks absorbed only in the visible region at longer wavelengths (461-463 nm), characteristic of all*-trans* isomers (O’neil and Schwartz, 1992; Seel et al., 2020).

**Figure 3.**
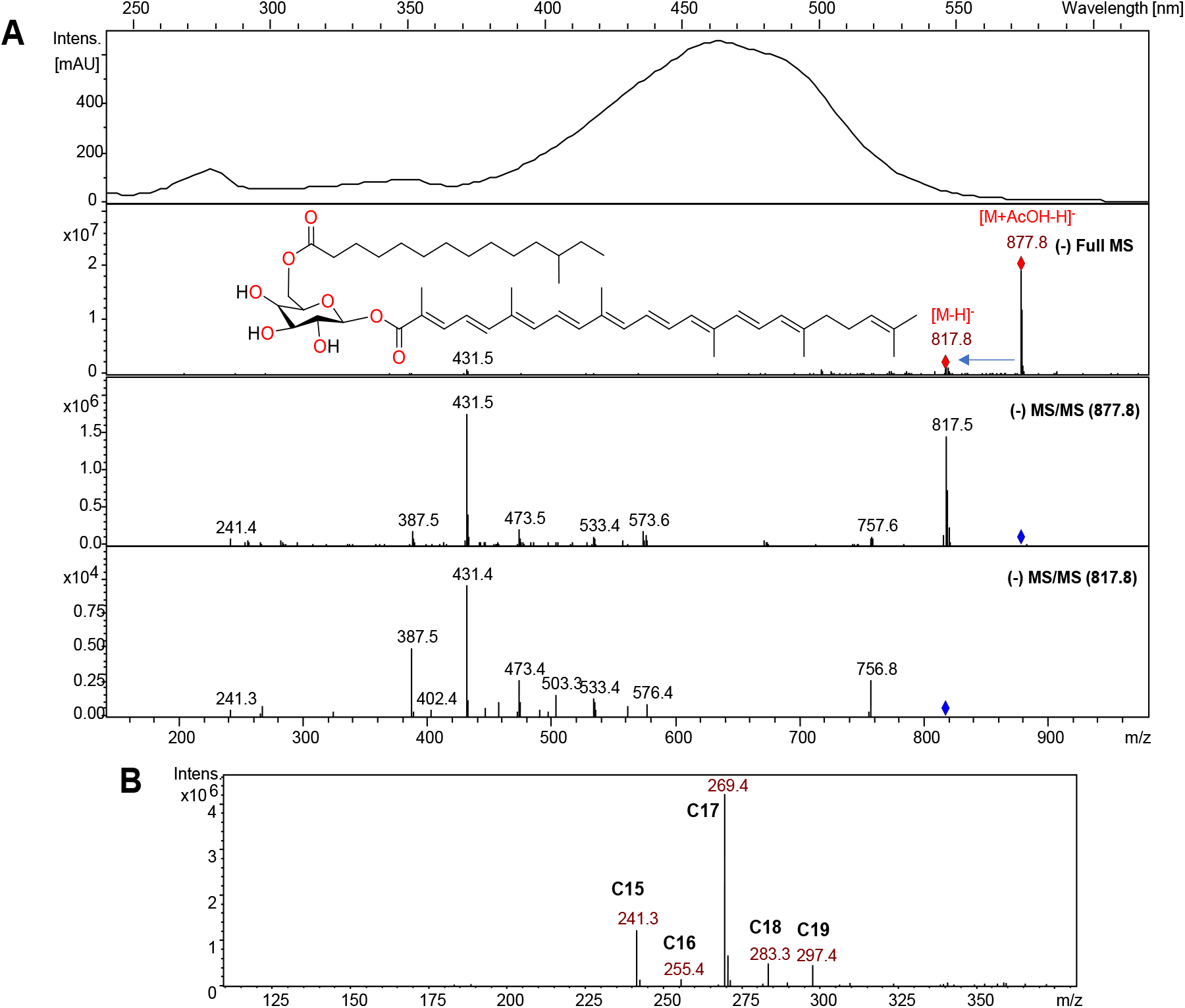
Spectrum of STX and free fatty acids. (A) UV-vis, Full MS and MS/MS spectra of STX. (B) Full MS spectra of fatty acids.

Four hydrocarbon carotenes, including 4,4’-diaponeurosporene (4,4’-DNP, M17), 4,4’-diapo-ζ-carotene (4,4’-DZC, M18), 4,4’-diapophytofluene (4,4’-DPF, M19), and 4,4’-diapophytoene (4,4’-DPE, M20), were identified in APCI(+), showing pseudo-molecular ions at *m/z* 403.5 [C_30_H_42_+H]^+^, *m/z* 405.5 [C_30_H_44_+H]^+^, 407.5 [C_30_H_46_+H]^+^, and 409.5 [C_30_H_48_+H]^+^, respectively, (Figure S4) (Marshall and Wilmoth, 1981b; Pelz et al., 2005; Taylor’ and Bavies, 1983). In addition, drastic changes in UV-visible absorbances for these metabolites allowed confirming their presence. Operating in ESI(+), only 4,4’-DNP was detected at *m/z* 402.5 [C_30_H_42_]^+^, with MS/MS fragments at *m/z* 387 and 310, which correspond to loss of methyl and toluene molecule, respectively (Kim and Lee, 2012; Marshall and Wilmoth, 1981a; Pelz et al., 2005). The carotene 4,4’-diapolycopene (4,4’-DLP, M16) was identified in the SA147 strain at *m/z* 401.5 [C_30_H_40_+H]^+^ in APCI(+), whereas in ESI(+) this carotene was observed at *m/z* 400.5 [C_30_H_40_]^+^ and tandem mass analysis presented the same fragments of 4,4’-DNP (Table 1). The carotenoids 4,4’-DLP and 4,4’-DLPA have been previously reported (Kim and Lee, 2012). Another interesting aspect to highlight is the bathochromic shift in UV-visible spectra analysis of carotenoids involved in the STX biosynthesis. Interestingly, as the degree of unsaturation increases, wavelengths shift from the UV to visible region and the multiplicity of the bands is lost due to the greater conjugation of the final molecules (4,4’-DNPA and STX) (**Fig. 2c**). Thus, the main six carotenoids involved in the STX biosynthetic pathway from *S. aureus* (4,4’-DPE, 4,4’-DPF, 4,4’-DZC, 4,4’-DNP, 4,4’-DNPA y STX) are characterized herein.

Three metabolites with the highest retention time were identified as menaquinones MK7 (M21), MK8 (M22), and MK9 (M23), with pseudo-molecular ions at *m/z* 649.7 [C_46_H_64_O_2_+H]^+^, *m/z* 717.7 [C_51_H_72_O_2_+H]^+^, and *m/z* 785.8 [C_56_H_80_O_2_+H]^+^, respectively, as observed in APCI(+) and in ESI(+) mode. As can be observed in table 1 and figure S5, menaquinones showed characteristic fragmentation ions at *m/z* 227 and *m/z* 187, and absorbances at (247 nm, 269 nm, and 329 nm), in accordance with data reported in literatura for *S. aureus* (Marshall and Wilmoth, 1981a; Taylor’ and Bavies, 1983; Wakeman et al., 2012). The presence of 4.4’-DNPA, STX and STX-homologues (STX-C13, STX-17, STX-C18 and STX-C19) was observed in the ATCC strain, in addition to the menaquinones MK7, MK8, and MK9.

The chromatographic elution obtained on the C18 column was according to the increasing polarity of the carotenoids, and provided a good resolution of *cis*/*trans* carotenoid isomers, typically reported for a C30 column (Amorim-Carrilho et al., 2014; Saha et al., 2019). Nevertheless, although STX is expected to elute before 4,4’-DNPA for its sugar moiety the final elution observed is according to the degree of polarity, similar to previous reports (Kim and Lee, 2012). The C18 chromatographic profiles of SA144, SA145, and SA147 strains extracts obtained after 24-hour growth were comparatively evaluated. While SA144 strain presented a similar carotenoid profile to SA401, SA145 only showed the characteristic menaquinone (M24-M26), as expected for the inhibition of carotenoids biosynthesis in this strain. In turn, SA147 presented an increase in the peak areas between 11 and 15 minutes, attributable to the reactivation of *S. aureus* carotenoid synthesis. (**Fig. 4a**). On the other hand, employing the C30 column, a similar resolution for *cis*/*trans* isomers in STX peaks was obtained compared to C18 column; however, carotenes 4,4’-DNP, 4,4’-DPF, 4,4’-DPE and menaquinones (MK7, MK8, MK9) are more broadly distributed throughout the chromatogram, due to the increasing resolution capacity of C30 stationary phase for isoprenoid derivates (Amorim-Carrilho et al., 2014; Saha et al., 2019). Nonetheless, the menaquinones mentioned above coelute with 4,4’-DNPA, STX or STX-homologues, therefore making more difficult the characterization (**Fig. 4b**).

**Figure 4.**
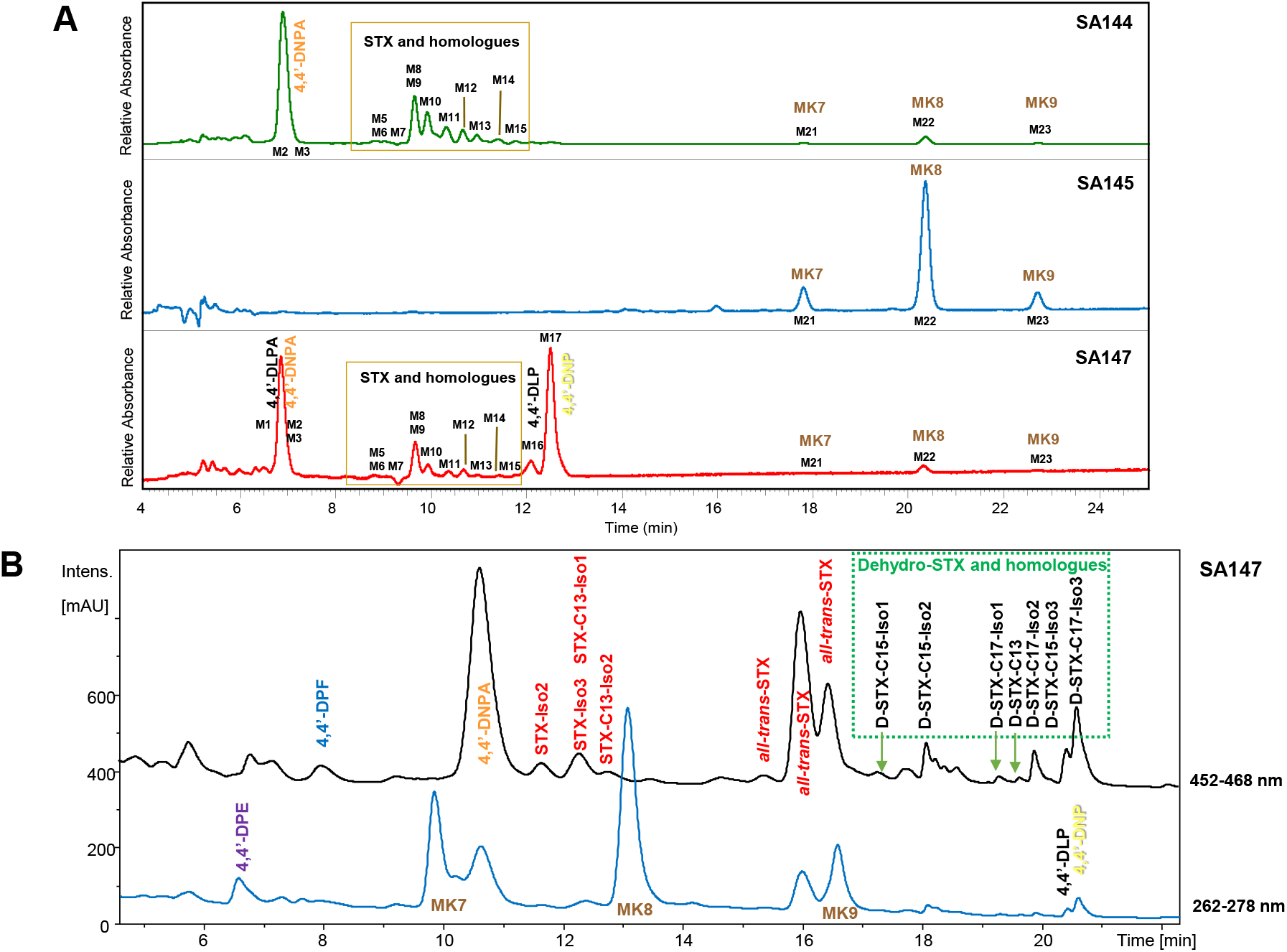
HPLC-MS analysis of *S. aureus* cells extracts in C18 and C30 column. (A) Chromatographic profile of SA144, SA145 and SA147 strains at 24 hours of culture in C18 column. (B) Chromatogram extracted in the visible region (322-338 nm and 422-436 nm) in C30 column.

Alternatively, the C30 column allowed the identification of new molecules in strain SA147, tentatively assigned as Dehydro-STX-C15, Dehydro-STX-C13 and Dehydro-STX-C17 (**Fig. 4b**), in addition to allow the confirmation of the carotenoid acid 4,4’-DLPA and hydrocarbon carotene 4,4’-DLP. Dehydro-STX-C13 (M30), Dehydro-STX (M26, M28 and M32), and Dehydro-STX-C17 (M29, M31 and M33) show [M+AcOH-H]^−^ adduct ions at *m/z* 848.0, *m/z* 875.7, and *m/z* 903.6, similarly to the adducts observed for STX and STX-homologues. MS/MS fragmentation in produced *m/z* 429 and *m/z* 385, related to the loss of CO_2_ (−44 amu) and typical masses of FA (**Fig. 5b**). In addition, Dehydro-STX and its homologues exhibited similar absorbances ((455), 483, 512 nm) to those reported for 4,4’-DLPA (Kim and Lee, 2012), a structurally related carotenoid (**Fig. 2**). Therefore, difference in two atomic mass units between fragments at *m/z* 431 (generated from STX-C13, STX and STX-C17) and *m/z* 429 (generated from Dehydro-STX-C13, Dehydro-STX and Dehydro-STX-C17) was attributed to an additional unsaturation in the carotenoid moiety. Thus, dehydro-STX homologues share structural similarity to their biosynthetic precursors 4,4’-DLP and 4,4’-DLPA also characterized in this work, as part of the alternative Dehydro-STX biosynthetic pathway illustrated in Fig. 1 (right side).

**Figure 5.**
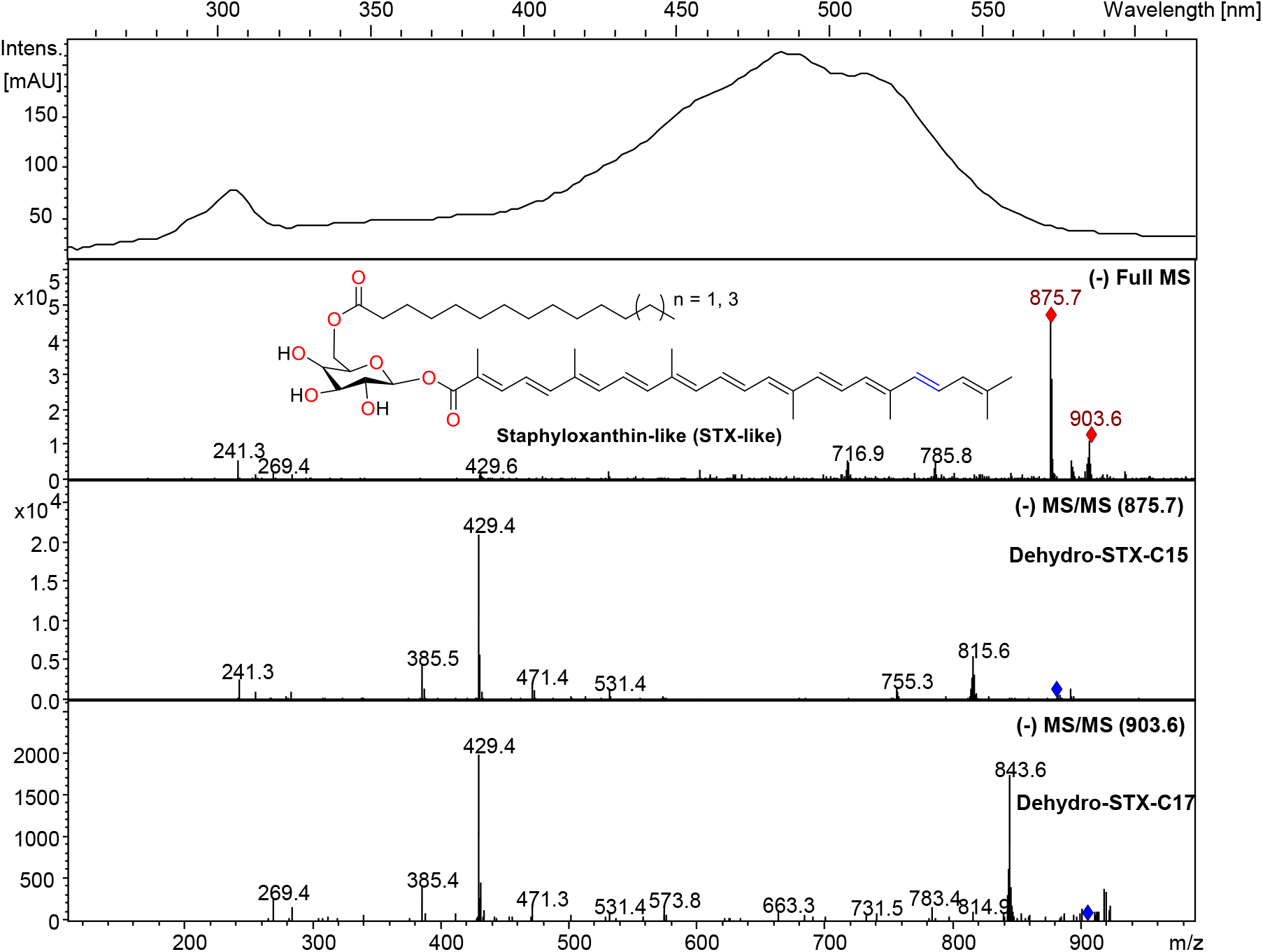
UV-vis, Full MS and MS/MS spectra for Dehydro-STX-homologues in SA147 strain.

Kinetic studies in *S. aureus* were performed by comparing the carotenoids profiles of SA401 *S. aureus* cell extracts obtained at different cell growth phases (8, 24 and 48 h), as illustrated in Fig. 6. Between exponential (8h) and stationary (24h) phases, a carotenes to xanthophylls interconversion can be clearly observed in the chromatographic profile (**Fig. 6**), which show the decrease or disappearance of carotenes: 4,4’-DPE, 4,4’-DPF, 4,4’-DZC, 4,4’-DNP, whereas the xanthophylls 4,4’-DNPA and STX (including STX-homologues) increase.

**Fig. 6.**
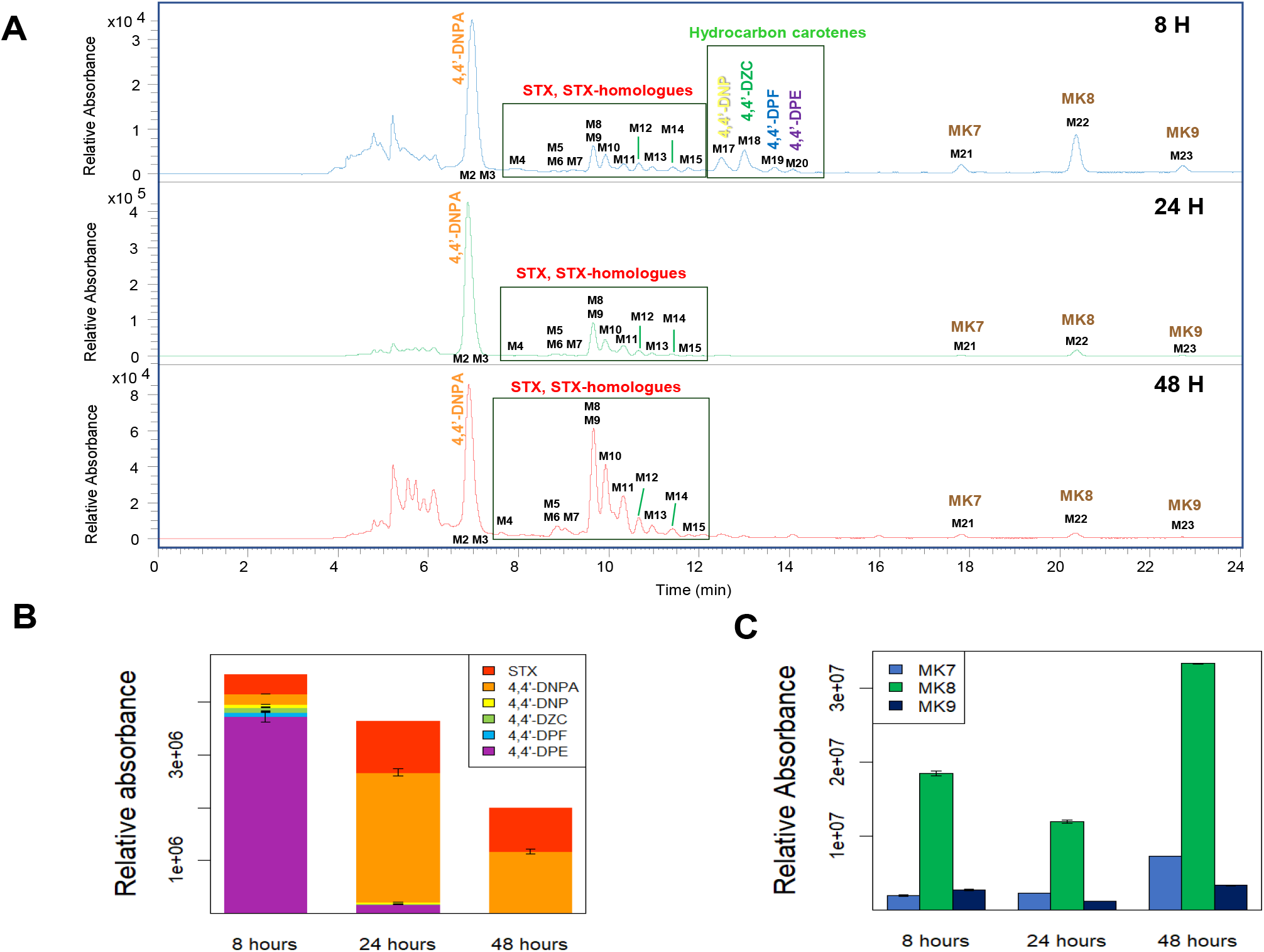
Carotenoids and menaquinones analysis from *S. aureus* cells at 8, 24, and 48 hours. (A) Comparative chromatograms of carotenoids at three growing times. (B) Metabolic changes in carotenes and xanthophylls composition at three growing times. (C) Metabolic changes in menaquinones composition at three growing times.

In addition, greater similarity between both chromatographic profiles (peaks between 7-12 min) at 24- and 48-hours of growth was observed (**Fig. 6a, 6b).** However, the persistence o 4,4’-DNPA in the two evaluated stationary states reflects that the increase in time does not lead to the exclusive presence of the final metabolite of this biosynthetic pathway: STX or STX-homologues. Besides, a variation in the relative areas of menaquinones is observed, displaying a fluctuation between the three growth times, show an increase in these metabolites at 48 h. (**Fig. 6c**). Additionally, relative abundance of menaquinones are more alike between 8 and 48 h compared to 24 h (**Fig. 6c**), which could be attributed to the fact that in the extreme phases of growth *S. aureus* prioritizes the synthesis of molecules that help it survive, since menaquinones are associated with cellular respiration, while carotenes and xanthophylls are secondary metabolites that have no direct function in survival unless there is stress external to the cells.

### 3.3. Membrane biophysical properties assessment

To understand the influence of carotenoids on the biophysical behavior of *S. aureus* membranes *in vivo*, wild type (SA144), knock-out (SA145), and regenerated (SA147) strains were comparatively evaluated by FTIR. Thus, the CH_2_ symmetric stretch band was analyzed between 2860 to 2840 cm^−1^ as a function of temperature changes from 5 to 50 °C, as shown in **Fig. 7**. Increasing wavenumber values can be related to changes in membrane lipid packing since the CH_2_ stretch indicates the level of trans/gauch isomerization in the acyl chains. A lower wavenumber indicates a high number of trans isomers that are associated to a straight acyl chain, which result in higher lipid packing. An increase in the wavenumber indicates an increase in gauch rotomeres related to a more disordered acyl chain in the phospholipid components of the membrane and an increase in lipid spacing. Lower temperatures favor the all trans configuration in lipids. Pure saturated phospholipids are characterized by cooperative first-order chain melting events that occur at specific melting temperatures. These are well documented transitions from a more tightly packed gel phase (L_β_) to a liquid crystalline phase (Lα). In pure phospholipid species such as 1,2-dipalmitoyl-sn-glycero-3-phosphocholine (DPPC) with Tm = 41 °C, 1,2-dimyristoyl-sn-glycero-3-phosphocholine (DMPC) at 24 °C, and several saturated Phosphatidyl Glycerol (PG) lipids characteristic of *S. aureus*. Although the more complex composition of bacterial membranes reduces the cooperativity of these melting events. thermotropic transition of the CH_2_ stretch vibration have been reported for *S. aureus* around 15 °C involving the cooperative melting of PG lipids contained in the plasma membrane of *S. aureus* (Ocampo et al., 2010; Scherber et al., 2009; Schultz and Naumann, 1991). Figure 7a shows cooperative melting events for SA144, SA145, and SA147 indicated by thermotropic shifts in the CH_2_ stretch vibration. The absence of carotenoid synthesis in strain SA145 results in an overall increase in the CH_2_ stretch wavenumber for temperatures above the phase transition temperature, and in particular at the growth temperature of *S. aureus* (Fig. 7b). This is direct indication of decreased chain order in the phospholipid acyl chains in the absence of carotenoids, and is consistent with previous studies using DPH and Laurdan (Perez-Lopez et al., 2019). When carotenoid synthesis is reestablished (SA147), the wavenumber drops. Indicating increased lipid packing. The results support the importance of carotenoids as regulators of lipid packing in *S. aureus* membranes (Mishra et al., 2011; Perez-Lopez et al., 2019; Sen et al., 2016; Tiwari et al., 2018).

**Fig. 7.**
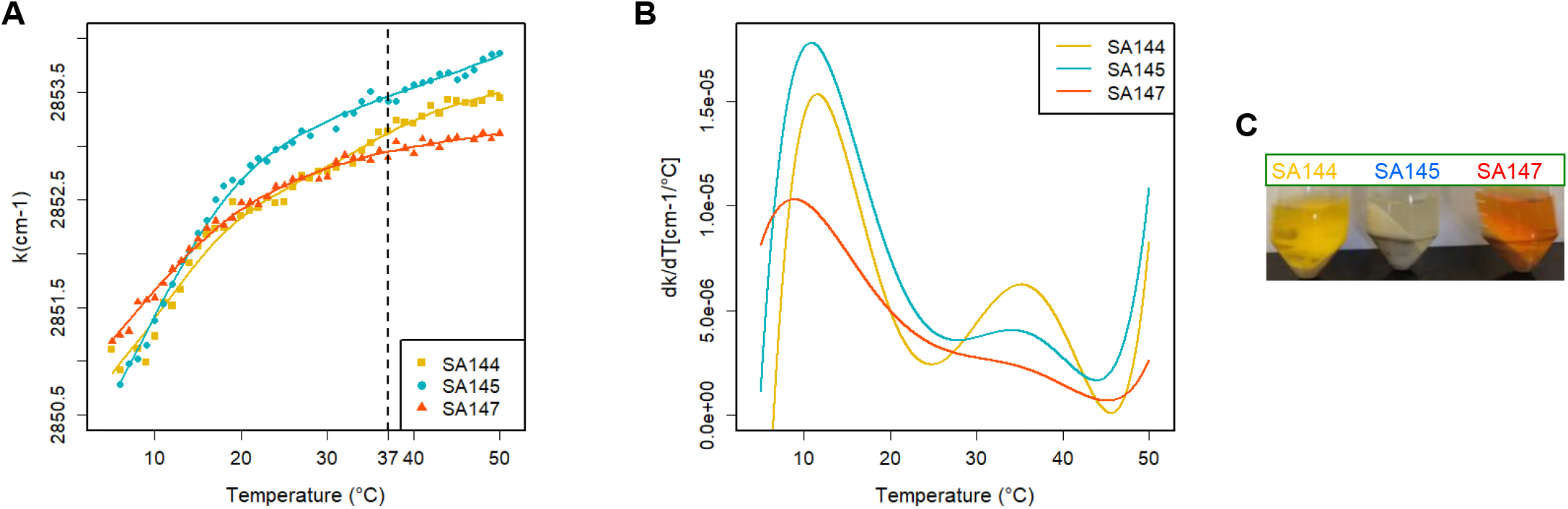
(A) Thermotropic phase behavior of the CH_2_ asymmetric stretch for *S. aureus in vivo* characteristic of the acyl chains in the membrane phospholipids in *S. aureus* for the native (SA144), CrtN knockout (SA145), and CrtN knockout incorporated with a plasmid containing CrtN (SA147) strains. (B) First derivative of the data obtained in (a) used to indicate the position of the main melting event and the cooperativity of the transition. (C) Different colors observed for extracts of SA144, SA145 and SA147.

The first derivative of the measurements in FTIR (**Fig. 7b)** indicate thermal events where the slope of the thermotropic curve is accentuated (See Fig 7a). These cooperative events reflect a change in the gauch/trans rotomer ratio for the acyl chain, indicating a clear change in the phospholipid packing behavior. The different packing levels of the membrane of *S. aureus* have been correlated to changes in the resistance of the membrane to antimicrobial agents (Bali et al., 2009; Mishra et al., 2011; Ocampo et al., 2010; Perez-Lopez et al., 2019). Strain SA144 shows two cooperative melting events appearing at around 10.5 and 33.0 °C. Strain SA145 exhibits a more accentuated change in the transition at around 11°C indicating that carotenoids tend to smooth the difference between the gel-like phase and the liquid-crystalline phase. This has been observed in model lipid systems in the presence of carotenoid extracts from *S. aureus* with the use of fluorescent probes and is confirmed here *in vivo* by FTIR (Perez-Lopez et al., 2019). This effect on the cooperativity of the chain melting event is similar to that observed in the presence of cholesterol in model systems and live cells (Bali et al., 2009). As carotenoid synthesis is reestablished (SA147) we observe a reduction in the cooperativity of the chain melting event (Fig. 7b SA147). In conclusion, Fig. 7 clearly indicates that carotenoids are regulators of membrane lipid packing in the *in vivo* system. at the growth temperature, and it is interesting to note that the small transition event that occurs around 35°C in the SA144 strain, also reported for the SA401 strain in a previous study (Ocampo et al., 2010), vanishes for SA145 and SA147. This small transition at a high temperature close to the growth temperature must be investigated further to be identified.

## 4. Discussion

The profiling analysis of *S. aureus* extracts obtained at different growth times led to the identification of six carotenoids belonging to the STX biosynthetic pathway. The six carotenoids (4,4’-DPE, 4,4’-DPF, 4,4’-DZC, 4,4’-DNP, 4,4’-DNPA and STX (including STX-homologues)) could be identified in the bacteria at 8 hours of cell culture. The relative abundance of these precursor compounds at 8 hours comparted with 24 or 48 hours reflects the reactivation of the STX biosynthesis route, which was shown to be downregulated in the early exponential phase (Perez-Lopez et al., 2019). In fact, the precursor 4,4’-DPE exhibited the highest abundance, (**Fig. 6b**) in accordance to the results reported by Wieland *et al.* (Wieland et al., 1994), who found a 50% lower concentration of STX compared with 4.4’-DNP after 12 hours of growth. The lower presence of xanthophylls at the initial growth stage is in line with proteomic studies of *S. aureus,* which also observed significant differences in protein expression levels between resuscitating and freezing survived cells (Suo et al., 2018). The metabolic change between the relative abundances of metabolites observed in the exponential state (8h) and those observed in the stationary states (24 h and 48 h), suggests that carotenoid biosynthesis reached the highest level of xanthophyll production late in the late exponential phase (Fig. 6b), reaching maturation of STX synthesis in the early stationary phase. The greater similarity in the proportion of metabolites when comparing the phase stationary profiles suggests that after reaching stationary phase *S. aureus* has a stable level of STX and 4,4’-DNPA. It is interesting to note that carotenoid acid 4,4’-DNPA remains stable for all growth states. Lower abundance of xanthophylls in general can be observed at the latter growth phase (48h) compared to the 24h cell culture (**Fig. 6b**), which can be explained by the depletion of nutrients in the LB culture medium (not renewed) in late stationary phase. These results are in good agreement with that reported by Wieland *et al.* In *S. aureus*, which indicate a greater amount of STX at 24 hours compared to 36 hours of cell growth (Wieland et al., 1994). Also, the observed increase in menaquinones content at 48 hours might be due to a stress for lack of nutrients (medium LB) on the microbiological system (**Fig. 6c**), that led to an increase in the production of these molecules associated with the respiration of the bacteria (Kurosu and Begari, 2010; Wakeman et al., 2012).

The reported structural diversity in fatty acid chains for *S. aureus* grown in LB broth, show a high concentration (77.2%) of branched-chain fatty acids (BCFAs), whereas straight-chain fatty acids (SCFAs) account for 22.8% (Sen et al., 2016), ranging from a C15 to C20 chain length. In this line, our characterization reveals the presence of a wide diversity of STX-analogues bonded to C13, C14, C16, C18, C19 and C20 fatty acid chains. Demonstrating that the acylation of glycosyl-4,4-diaponeurosporenoate mediated by CrtO enzyme is not exclusive to C15 and C17 fatty acids, as initially reported (Marshall and Wilmoth, 1981a), thus, C13 to C20 fatty acid chains that have been previously reported on the SA401 strain (Perez-Lopez et al., 2019) and other wild type *S. aureus* strains (Braungardt and Singh, 2019; Sen et al., 2016) were observed bonded to Glu-4.4’-DNPA core. The broad specificity of the acyltransferase CrtO was also reported in *E. coli* mutants with the formation of STX analogues derivatives that include C14 and C16 fatty acids and other fatty acids of different lengths of not yet characterized (Kim and Lee, 2012). Furthermore, the presence of a Staphyloxanthin derivate with three additional units of sugars, a molecule associated with microdomains generated by *S. aureus,* was recently indicated (García-Fernández et al., 2017), confirming the diversity of homologues in the synthesis respect to this carotenoid with saccharolipid nature. STX analogues with such broad range of fatty acid chains, has not been reported in *S. aureus* grown in LB media. We believe that STX-analogues have higher proportion of BCFAs, according to previous reports (Perez-Lopez et al., 2019; Sen et al., 2016; Tiwari et al., 2018).

In this regard, previous reports indicate that CrtN knock-out strains harboring a plasmid containing CrtN, generated carotenoid extracts of red color, associated with the presence of additional alternative metabolites (Umeno et al., 2005). Thus, carotenoid extracts in wild type *S. aureus* strains are orange color, whereas *S. aureus* strains where CrtN has been reincorporated through a plasmid are reddish, with products which are alternative to the main carotenoids biosynthetic pathway (Kim and Lee, 2012). The regenerated strain SA147 presents additional desaturase activity that converts 4,4’-DNP into the red-colored 4,4’-DLP carotene, as observed in the extract of these cells (**Fig. 7c**), as well as 4,4’-DLPA due to the oxidation of the 4,4’-DLP. Similar visible region absorptions and characteristic *m/z* values suggest that this strain has the capacity to generate three compounds, here denominated Dehydro-STX and its homologues, through the alternate route indicated in the right part of figure 1a. The characterization of Dehydro-STX, Dehydro-STX-C13, and Dehydro-STX-C17 described herein pose a valuable contribution to the work previously reported (Kim and Lee, 2012). In addition, the presence of STX homologues in the ATCC strain indicates that variation in fatty acid chains is not exclusive to the other strains studied. However, the C15 fatty acid chain is still predominant in proportion to the other FAs in all strains.

In light of the results, it is worthy to highlight that the profiling methodology proposed in this work made it possible to characterize STX and Dehydro-STX with their respective homologues, by analyzing the extract instead of fractions obtained by TLC or OCC separations as reported in previous works (Kim and Lee, 2012; Marshall and Wilmoth, 1981a; Pelz et al., 2005). Thus, the profiling approach proposed in this work avoids overlooking minor compounds. Another remarkable analytical aspect is the *cis*/*trans* isomers resolution capacity shown by the C18 column, comparable to C30 columns, commonly used for carotenoids separation and for resolution of geometric isomers. These results add to the previously reported efficiency of C18 columns for the analysis of xanthophylls (Amorim-Carrilho et al., 2014; Saha et al., 2019). Besides, the resolution capacity demonstrated by the C18 could be explained as the greater interaction between this stationary phase and the analyte, that depends on the combination of hydrophobicity and dispersion forces, while theC30 relies exclusively on the hydrophobicity of the interaction with the carotenoids (Saha et al., 2019). This was also corroborated by the change in the order of elution of the carotenes, from the most non-polar to the most polar 4,4’-DPE, 4,4’-DPF, 4,4’-DNP, in accordance with the more hydrophobic character of the initial mobile phase. Also, the broad distribution of metabolites throughout the chromatogram showed by triacontyl (C30) stationary phase generates characterization issues, due to coelution of xanthophylls and menaquinones, as described in the results section. However, an outstanding aspect of the C30 column was its resolution capacity in the xanthophylls of the SA147 strain, as it allowed the characterization of the new Dehydro-STX include their homologues and allowed the completion of the alternate biosynthetic pathway described here.

Decrease in the membrane fluidity of *S. aureus* has been associated with the increase of STX content in the bacteria (Mishra et al., 2011; Perez-Lopez et al., 2019; Sen et al., 2016), hence the interest in the study of this secondary metabolite. However, many of these reports about the membrane biophysical behavior of *S. aureus* do not consider the intermediate species that may be present in the crude extract, mainly characterized by UV-vis spectrophotometry. In addition, previous studies on STX biosynthesis using mutant strains of *S. aureus*, *S. carnosus*, and *E. coli* (Kim and Lee, 2012; Pelz et al., 2005), showed the coexistence of intermediate species in the carotenoid extract. Thus, the results reported here on two wild-type strains of *S. aureus* (SA401 and SA144) and one mutant (SA147) allow us to confirm that this coexistence of two major species (STX or STX-homologues and 4,4’-DNPA) observes at 24- and 48-hour of growth is proper to the bacterium. This aspect should be considered when proposing model compositions to study biophysical aspects of *S. aureus* membranes, including studies related to the activity of antimicrobial peptides. Since in different reports, the inhibition of antimicrobial peptide activity is only attributed to the presence of STX and not to other carotenoid metabolites (Mishra et al., 2011; Sen et al., 2016). According to the above, we hypothesize that the precursor carotenoid 4,4’-DNPA is an intrinsic component of the *S. aureus* membrane, which is likely to play an important role in regulating membrane stiffness. This species does not contain the additional sugar group or acyl chain that is present in STX and should be treated as a free fatty acid with a highly rigid chain group inserted in the membrane.

Finally, FTIR data on *S. aureus* cell *in vivo* show two distinct results. First, the level of acyl chain order, measured as the proportion of gauche/trans rotometer in the acyl chains of the phospholipids, increases significantly in the presence of carotenoids in the high temperature range, which includes the growth temperature. This is consistent with previous studies using indirect methods for measuring acyl chain order and headgroup spacing using extrinsic fluorescent probes such as DPH and Laurdan (Perez-Lopez et al., 2019). Additionally, the FTIR results show that the cooperativity of the main melting event around 11°C does not present a significant shift in temperature when carotenoids are present. However, the level of cooperativity of this transition event, measured as the intensity of the first derivative of the thermograms, is greatly reduced when carotenoids are present. This behavior is consistent with that observed for the incorporation of cholesterol into mammalian cells, where the presence of cholesterol increases chain order in the liquid-crystalline phase, inducing the formation of a liquid-ordered phase (Bali et al., 2009; Gousset et al., 2002; Mannock et al., 2006). In addition, cholesterol has been shown to reduce the cooperativity of the gel to liquid-crystalline phase. This effect is related to the rigid and planar ring structure of the cholesterol molecule which forces the acyl chain of neighboring lipids in the liquid-crystalline phase to increase the proportion of trans rotamers (Bali et al., 2009; Mannock et al., 2006; Potrich et al., 2009). The rigid and extended structure of the tripenoids characteristic of STX in *S. aureus* appear to serve a similar function (Mishra et al., 2011). Recent, publications have presented evidence to indicate that STX homologues is involved in the formation of lipid domains (García-Fernández et al., 2017), and that these STX-enriched lipid domains play a role in methicillin resistance. This propensity to form lipid domains is likely related to the biophysical properties of the molecule and needs to be studied further to elucidate the mechanism by which these lipid domains are formed.

## 5. Conclusions

We employed a suitable LC-MS^n^ method for the analysis of the carotenoids present in *S. aureus* with minimum sample preparation of the extract. The joint spectral information allows the simultaneous analysis of carotenes, xanthophylls, and menaquinones from *S. aureus*. The tentative identification of 34 carotenoids and menaquinones produced by the microorganism was achieved and it is clear that STX is not the main component, even when the carotenoid composition is stabilized in the stationary phase, although it is responsible for the characteristic color of the bacteria. Also, the use of the ion trap (IT)-MS in this method allowed the complete identification of characteristic patterns of fragmentation of carotenoids, including new unreported molecules in knockout *S. aureus* strain incorporated with a CrtN containing plasmid. Results based on growth times show the sequential progression of metabolite precursors during late exponential phase (8 hours) leading towards a mature carotenoid profile of end products which includes 4,4’-DNPA and STX as the main components in the stationary phase (24 and 48 hour). These results reveal that in the interest of studying these most carotenoids of this microorganism, it is best to carry out its culture and extraction at 8 hours. Eventually, this method could lead to performing quantitative analysis of carotenoids in *S. aureus* and other microorganisms to identify intermediate species in different biosynthesis routes. In addition, the characterization of the melting temperatures in the fatty acid chains was achieved using FTIR and associated with increase acyl chain order in the presence of carotenoids and changes in the cooperativity of the membrane melting events.

## Acknowledgements

The authors wish to thank the Chemistry and Physics Departments and Research Fund of the Faculty of Sciences of the Universidad de los Andes for the financial support (INV-2019-86-1843). Also, to Ministerio de Ciencia, Tecnología e Innovación (MinCiencias) by National Fellowship No. 785 to Gerson-Dirceu López and the project EXT-2017-82-1779, and the grant No. 120480763040 of MinCiencias. The authors also thank the support from the AGL2017-89417-R project (Ministerio de Ciencia y Universidades).

