## Supplementary material for "Carotenogenesis of *Staphylococcus aureus*: new insights and impact on membrane biophysical properties": FigureS1

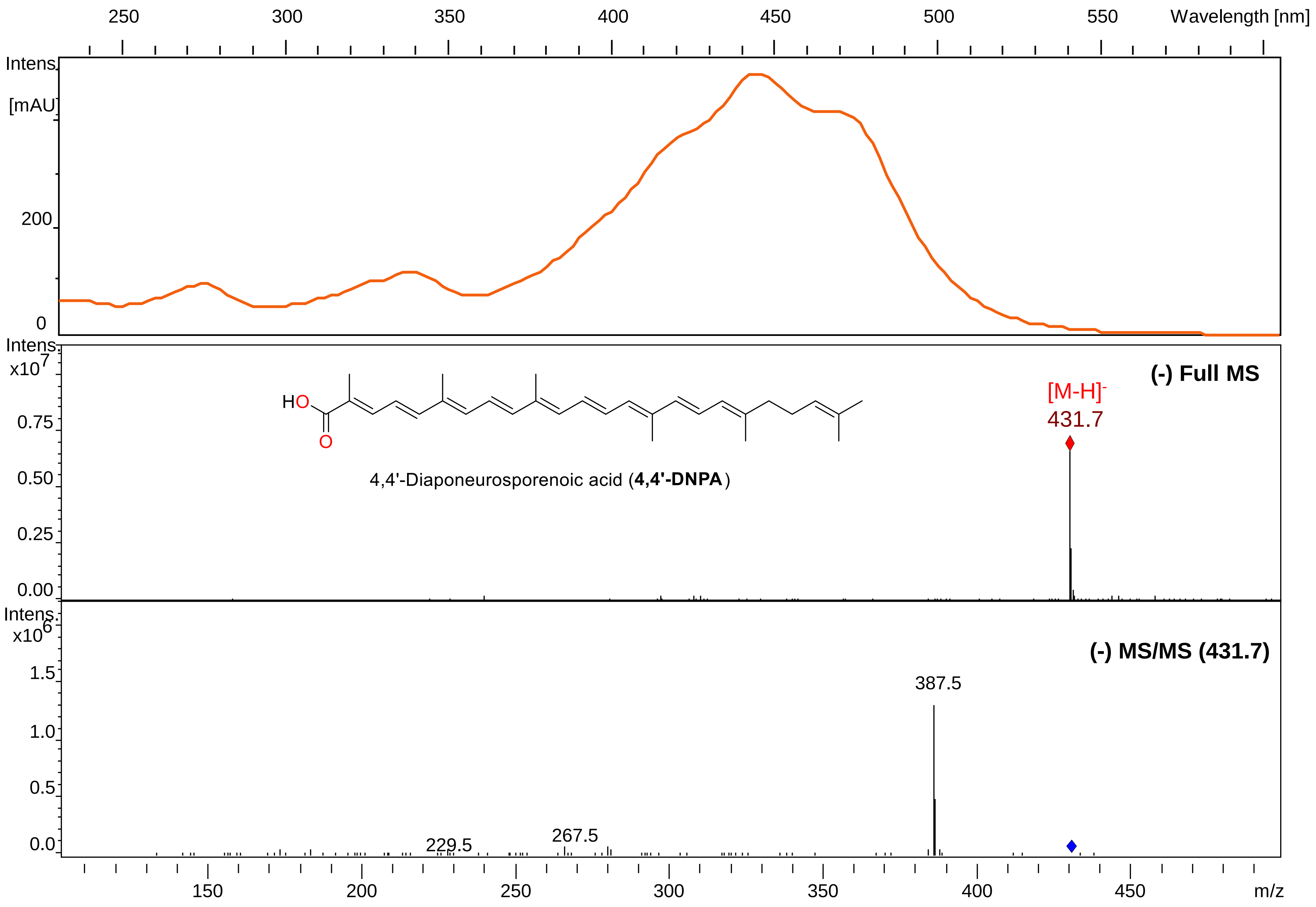


UV-visible, MS and MS/MS spectra for 4,4'-DNPA in APCI (-).

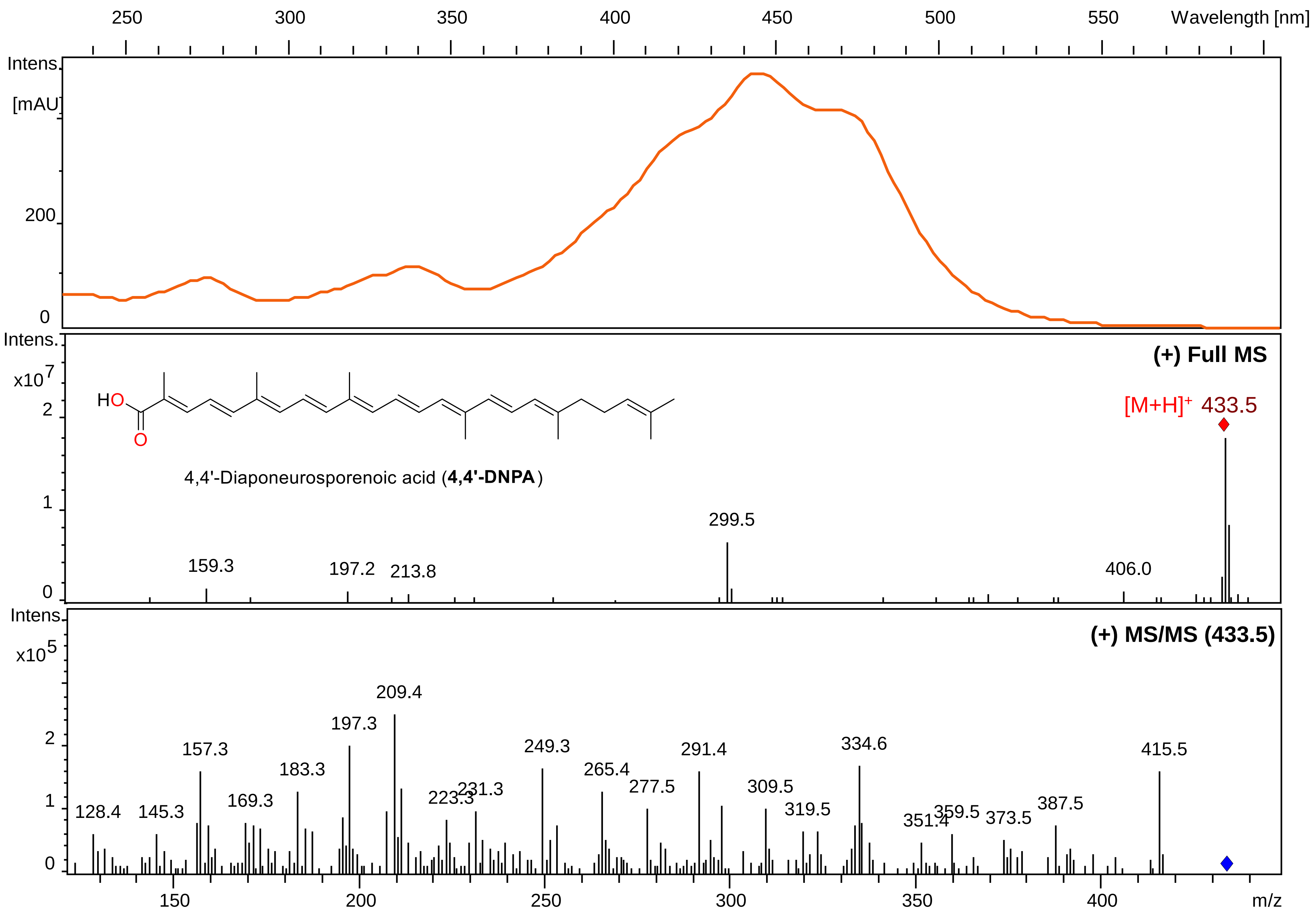


UV-visible, MS and MS/MS spectra for 4,4'-DNPA in APCI (+).

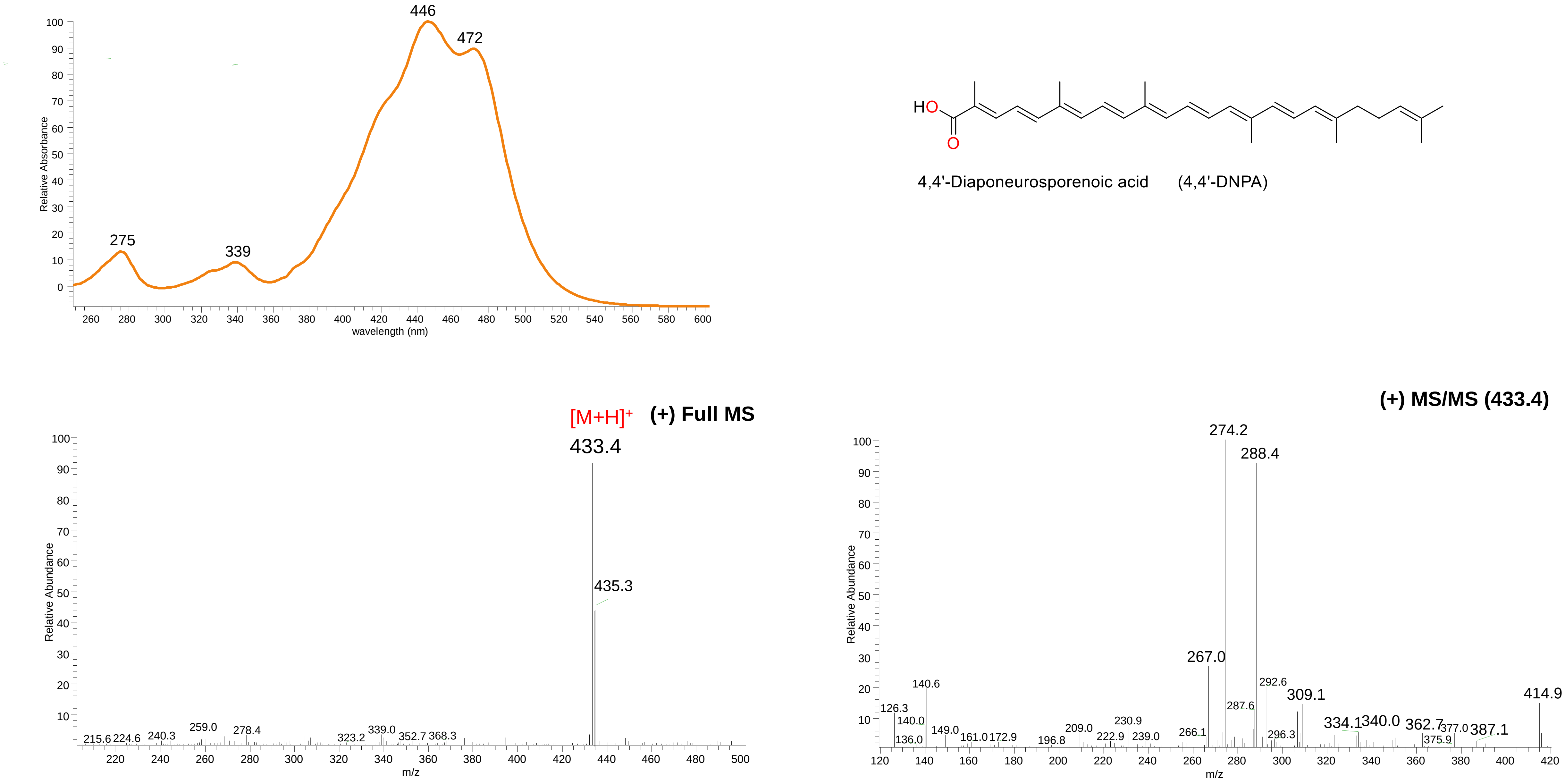


UV-visible, MS and MS/MS spectra for 4,4'-DNPA in ESI (+).

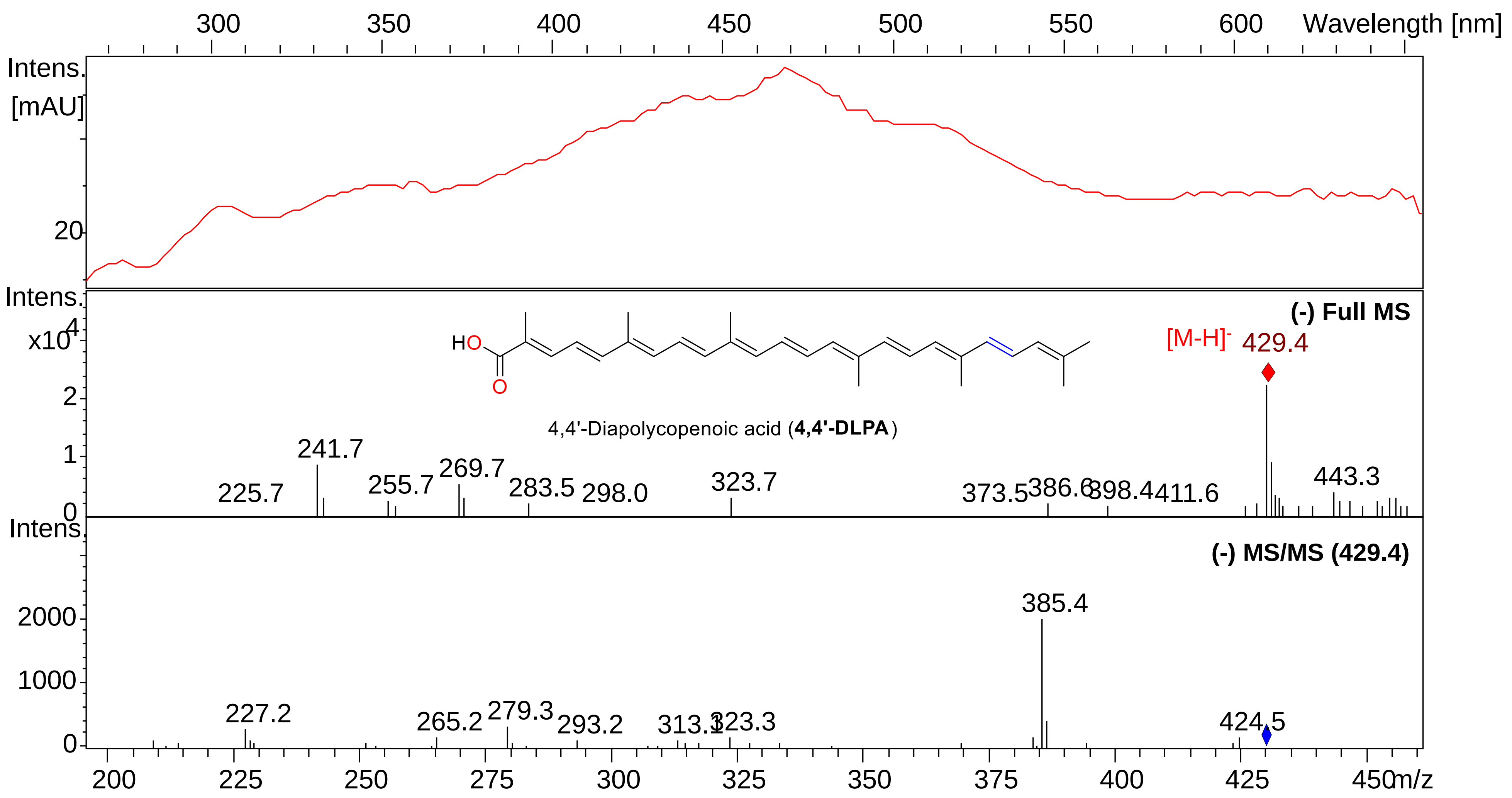


UV-visible, MS and MS/MS spectra for 4,4'-DLPA in APCI (-).
