## Supplementary material for "Carotenogenesis of *Staphylococcus aureus*: new insights and impact on membrane biophysical properties": FigureS2

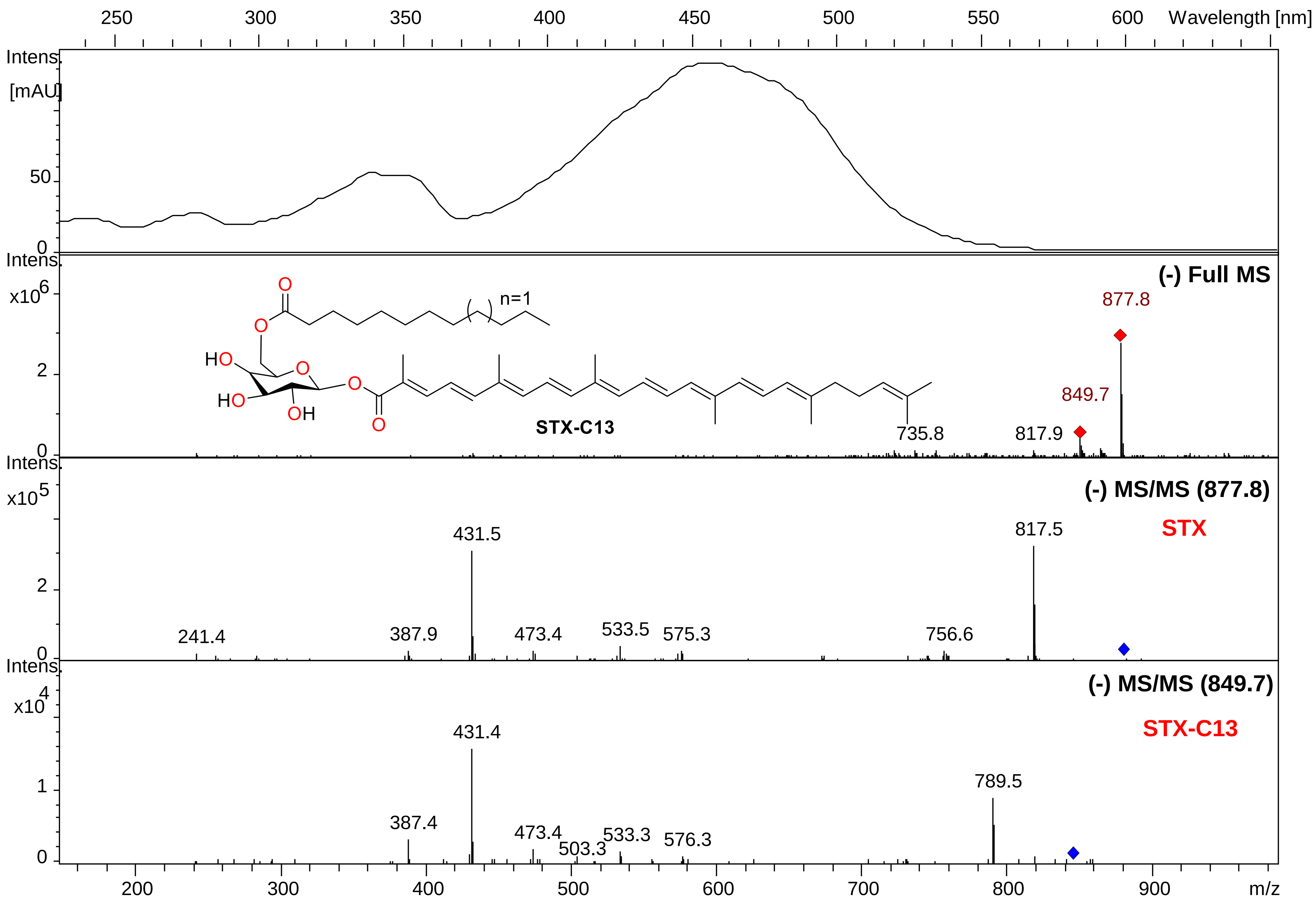


UV-visible, MS and MS/MS spectra for STX-C13 in APCI (-).

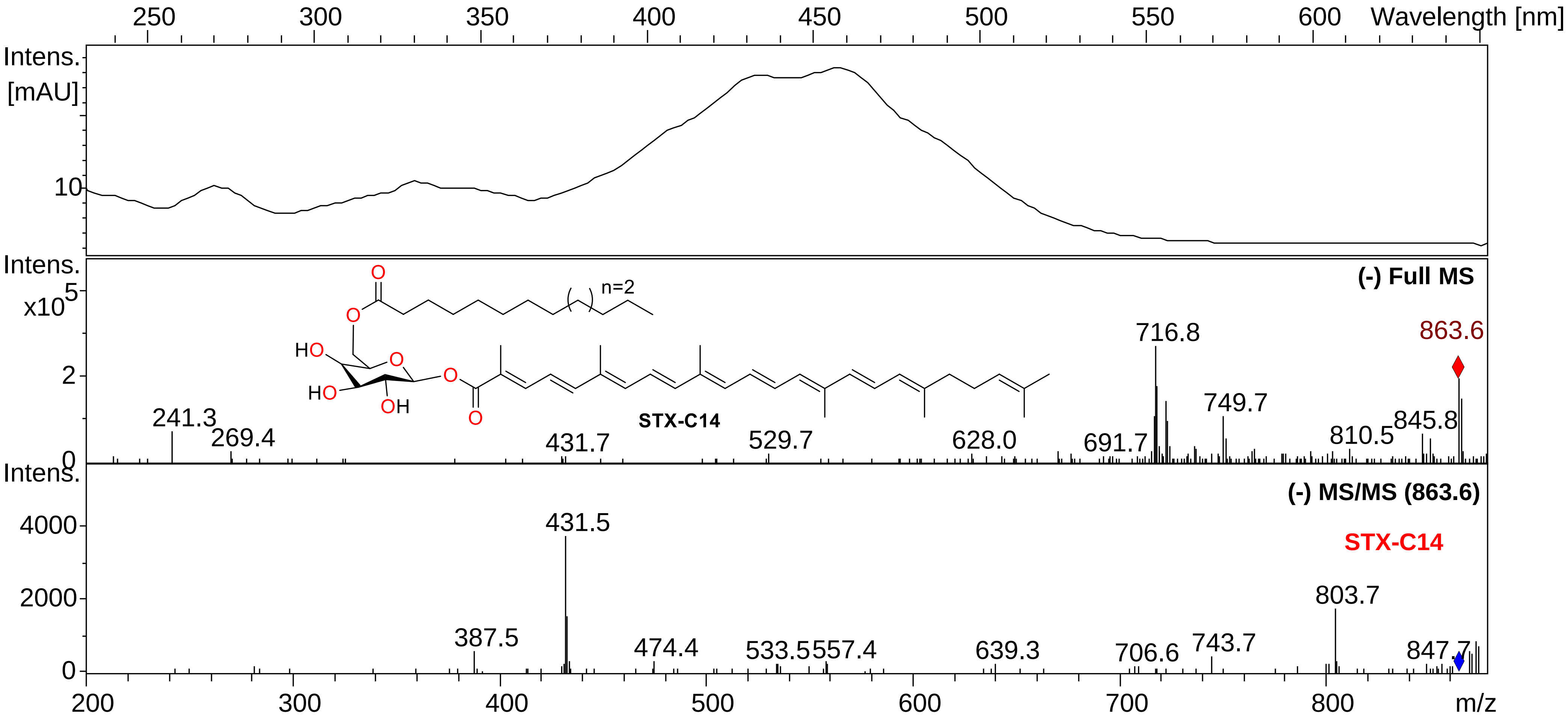


UV-visible, MS and MS/MS spectra for STX-C14 in APCI (-).

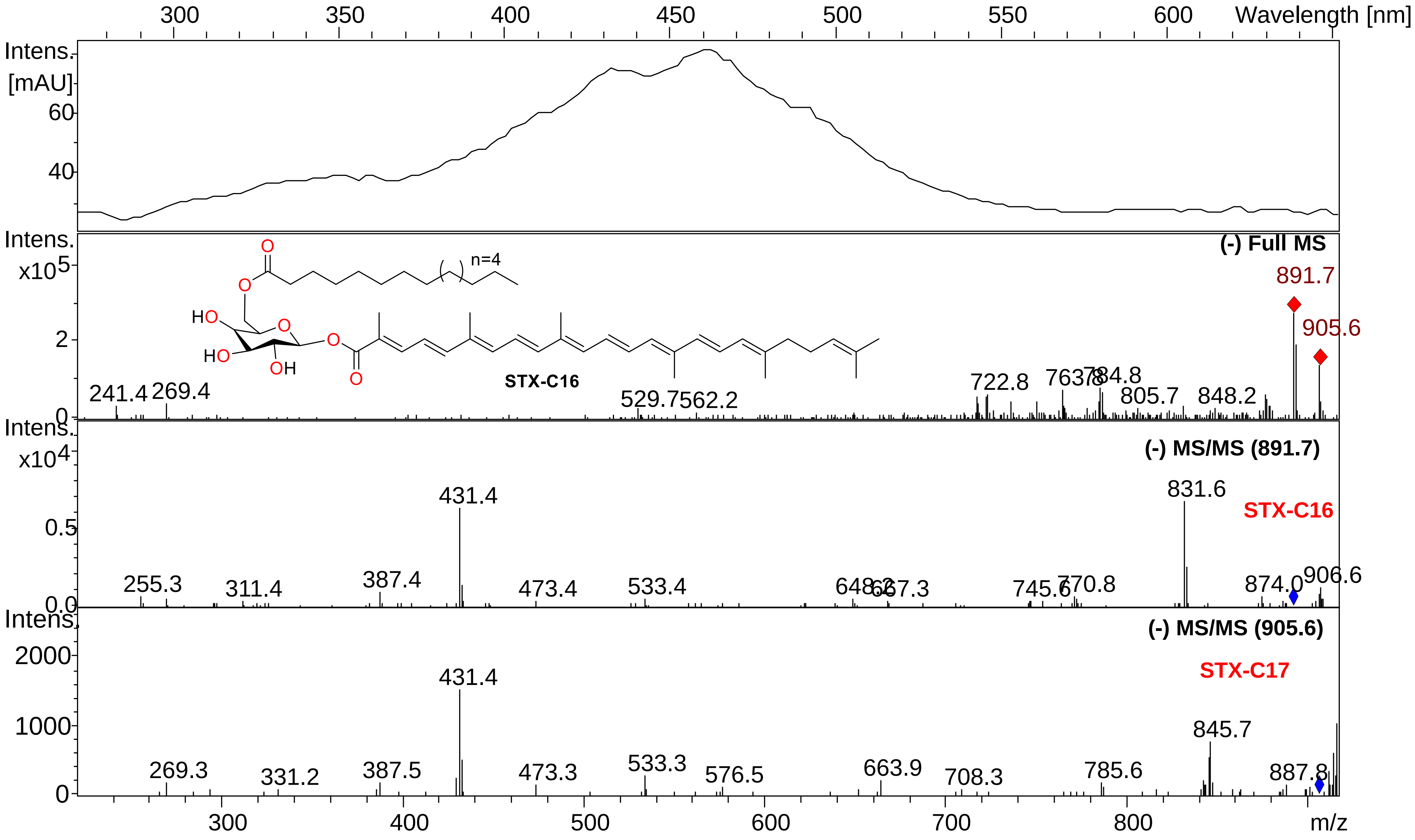


UV-visible, MS and MS/MS spectra for STX-C16 in APCI (-).

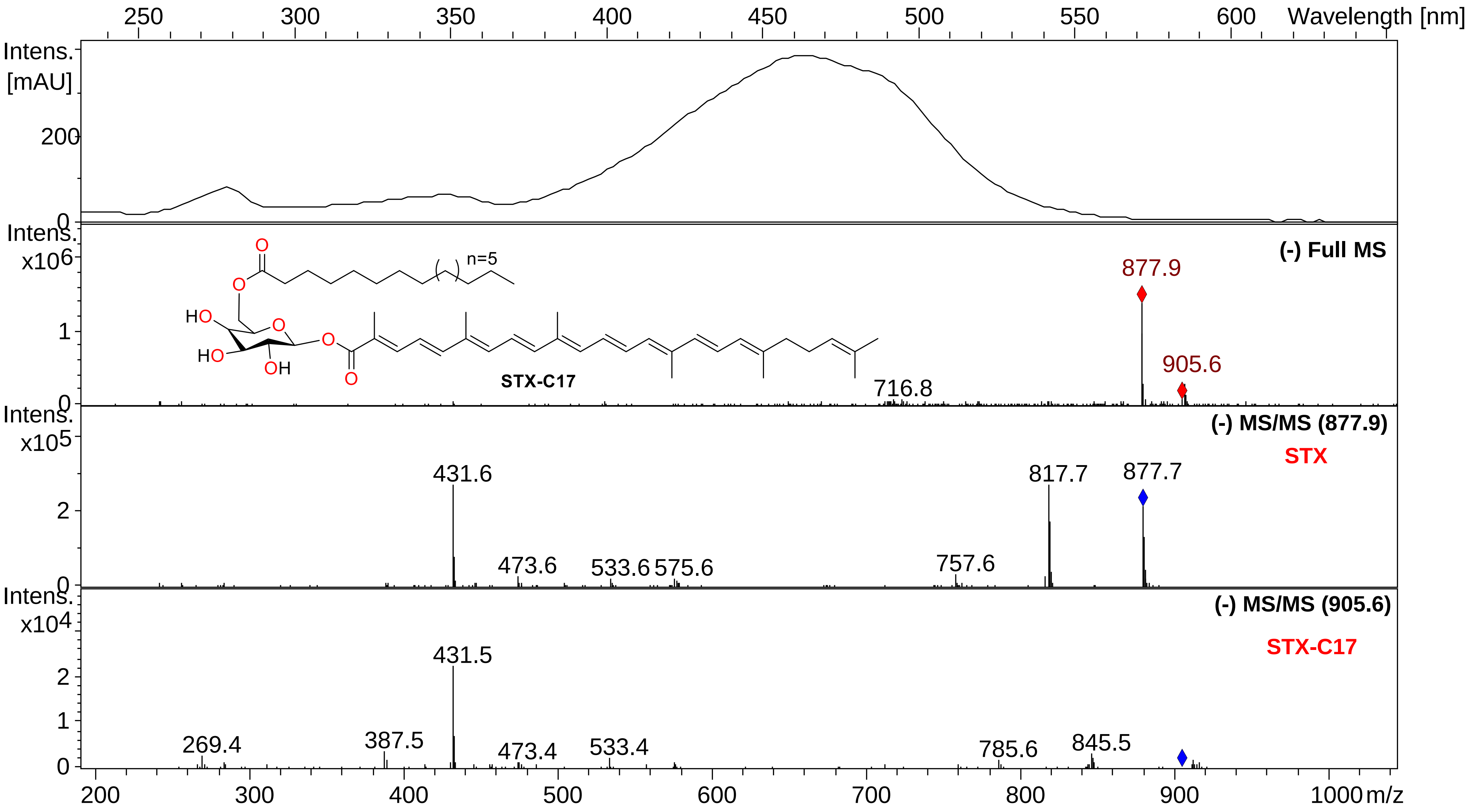


UV-visible, MS and MS/MS spectra for STX-C17 in APCI (-).

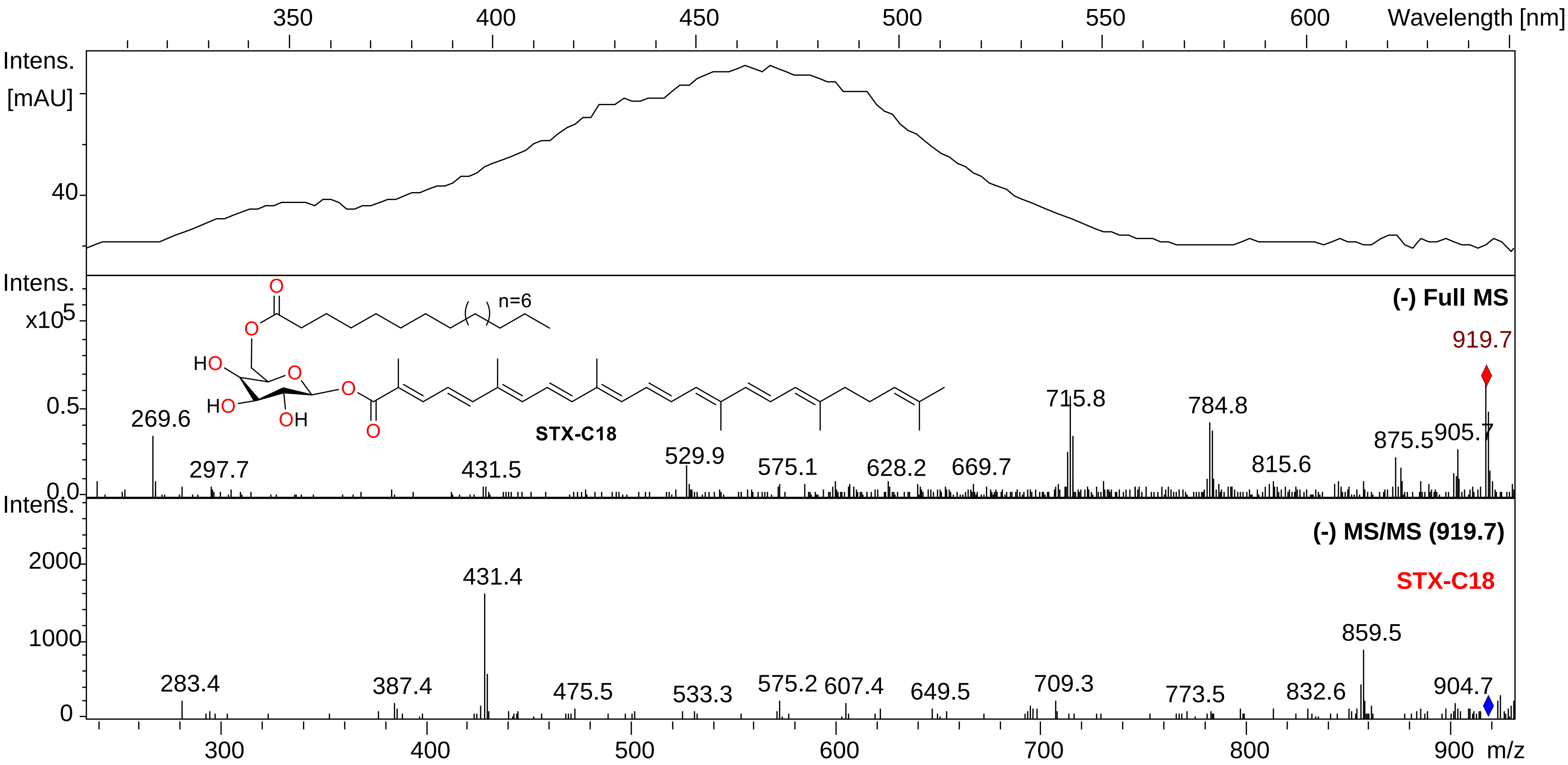


UV-visible, MS and MS/MS spectra for STX-C18 in APCI (-).

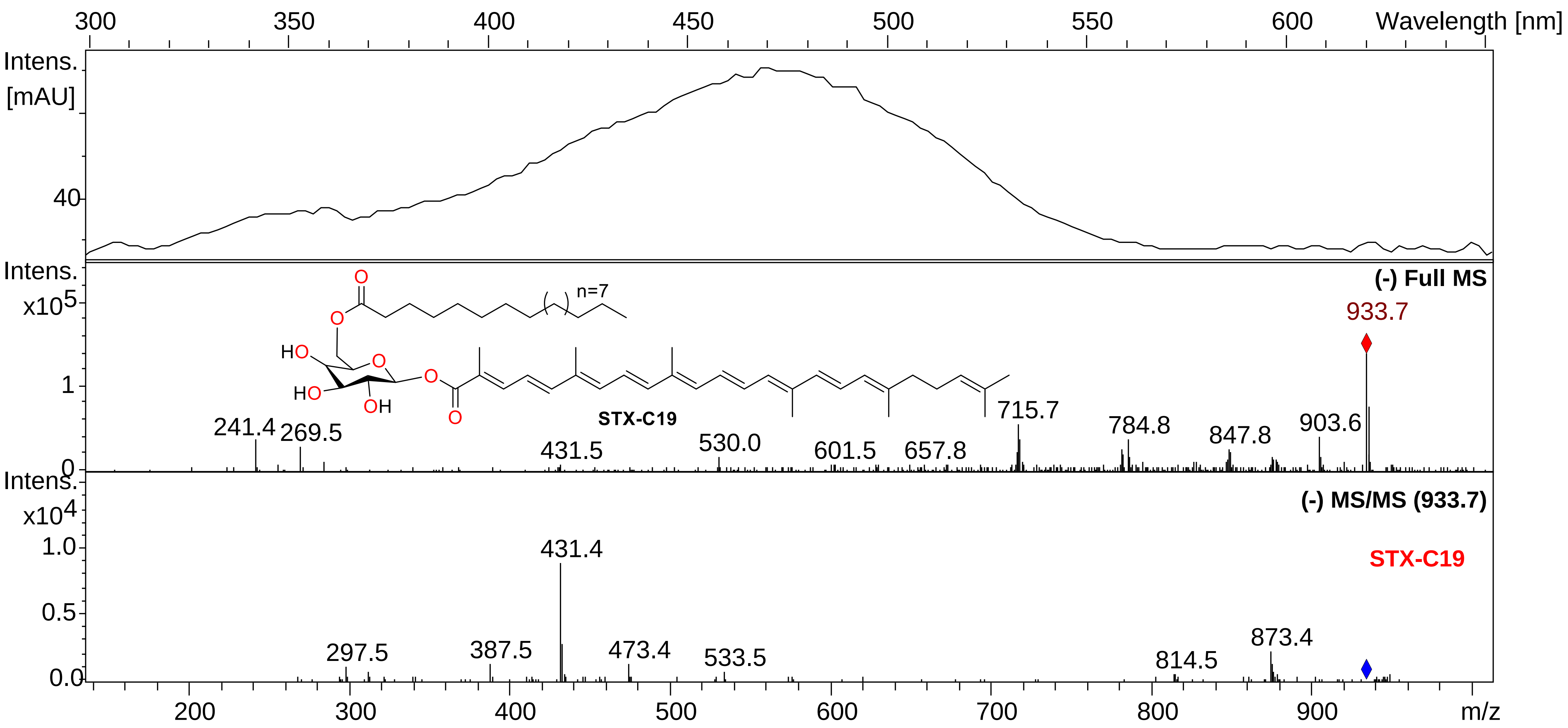


UV-visible, MS and MS/MS spectra for STX-C19 in APCI (-).

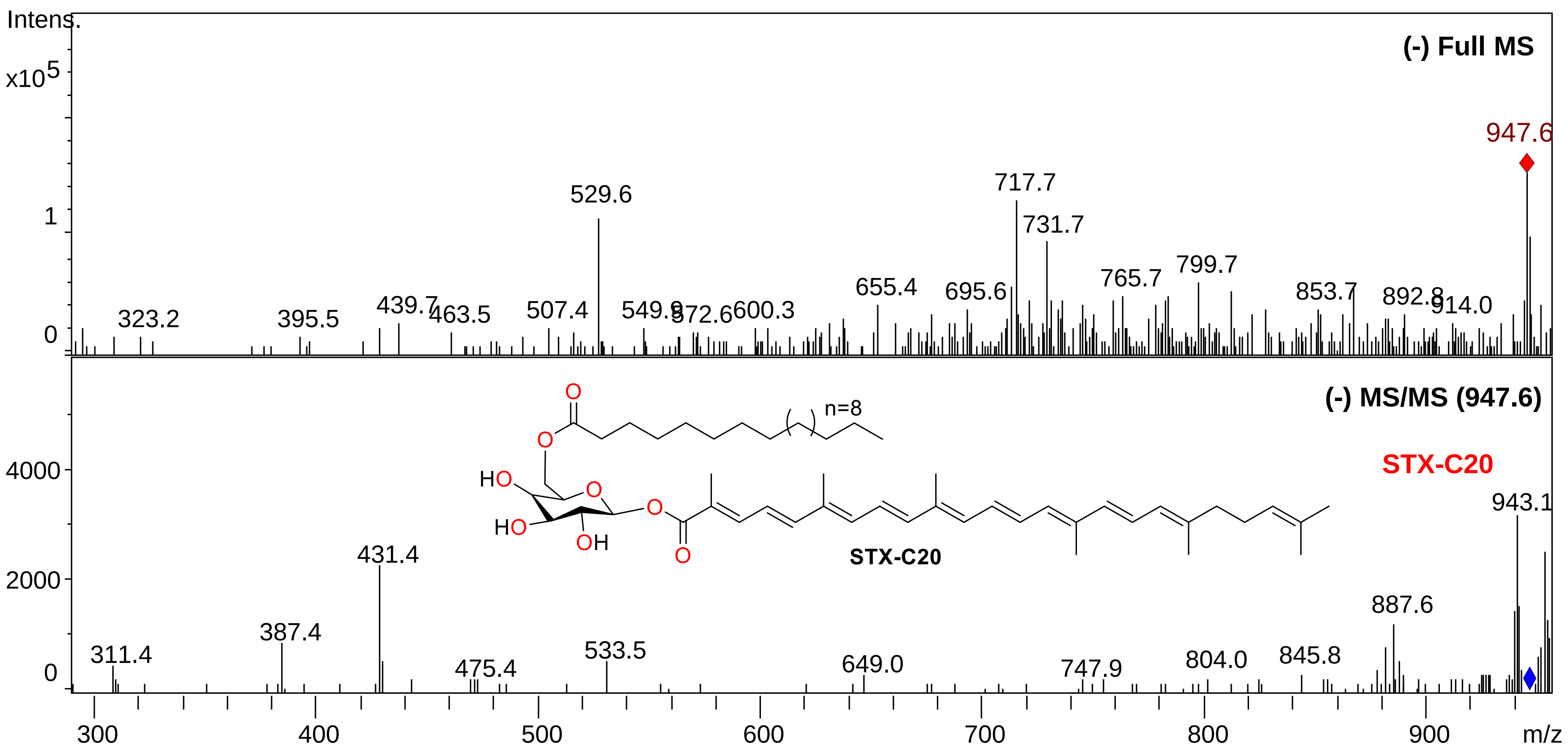


MS and MS/MS spectra for STX-C20 in APCI (-).
