## Supplementary figures and images for "Carotenogenesis of *Staphylococcus aureus*: new insights and impact on membrane biophysical properties"

### FigureS3

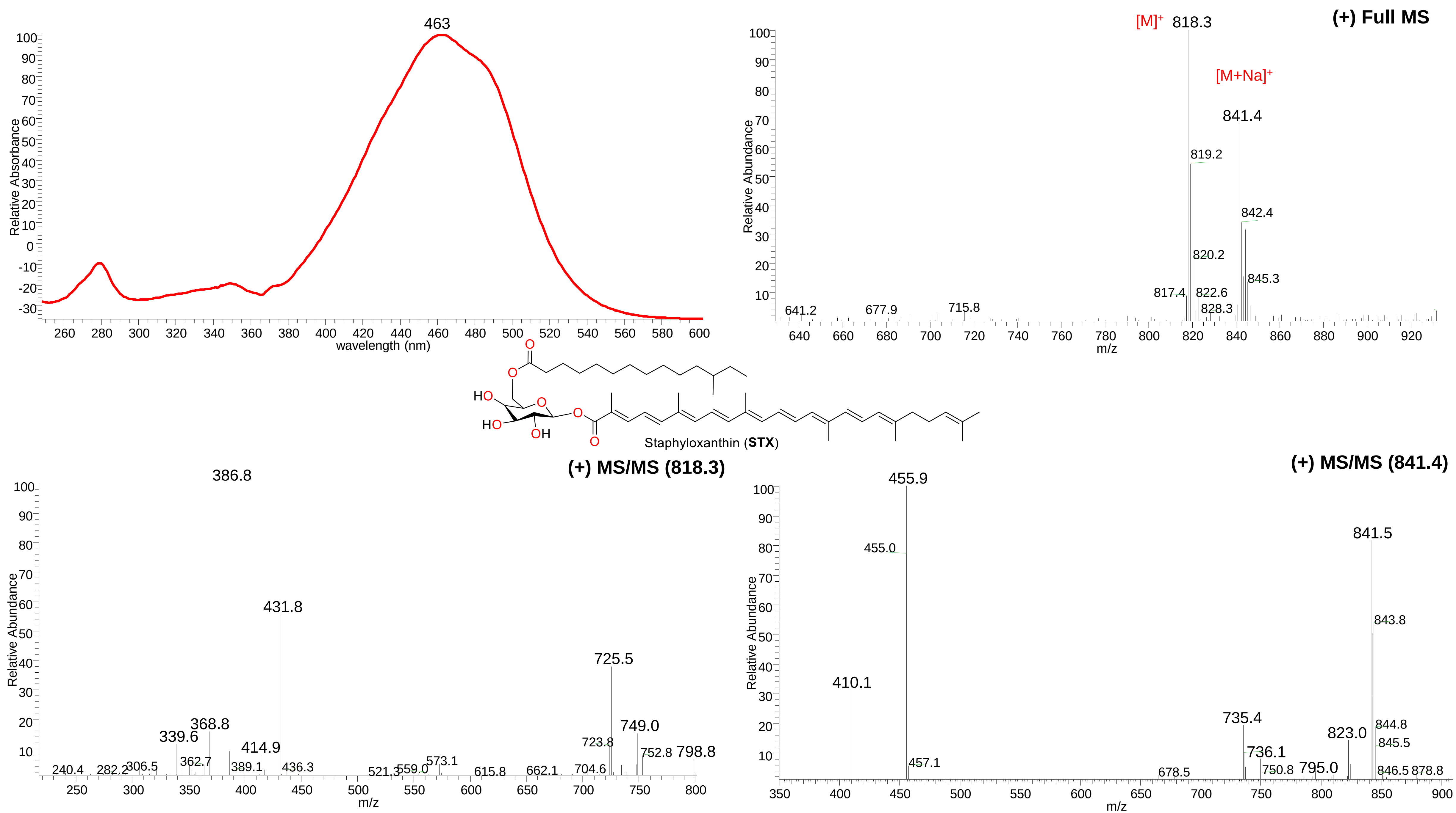


UV-visible, MS and MS/MS spectra for STX in ESI (+).
