## Supplementary material for "Carotenogenesis of *Staphylococcus aureus*: new insights and impact on membrane biophysical properties": FigureS4

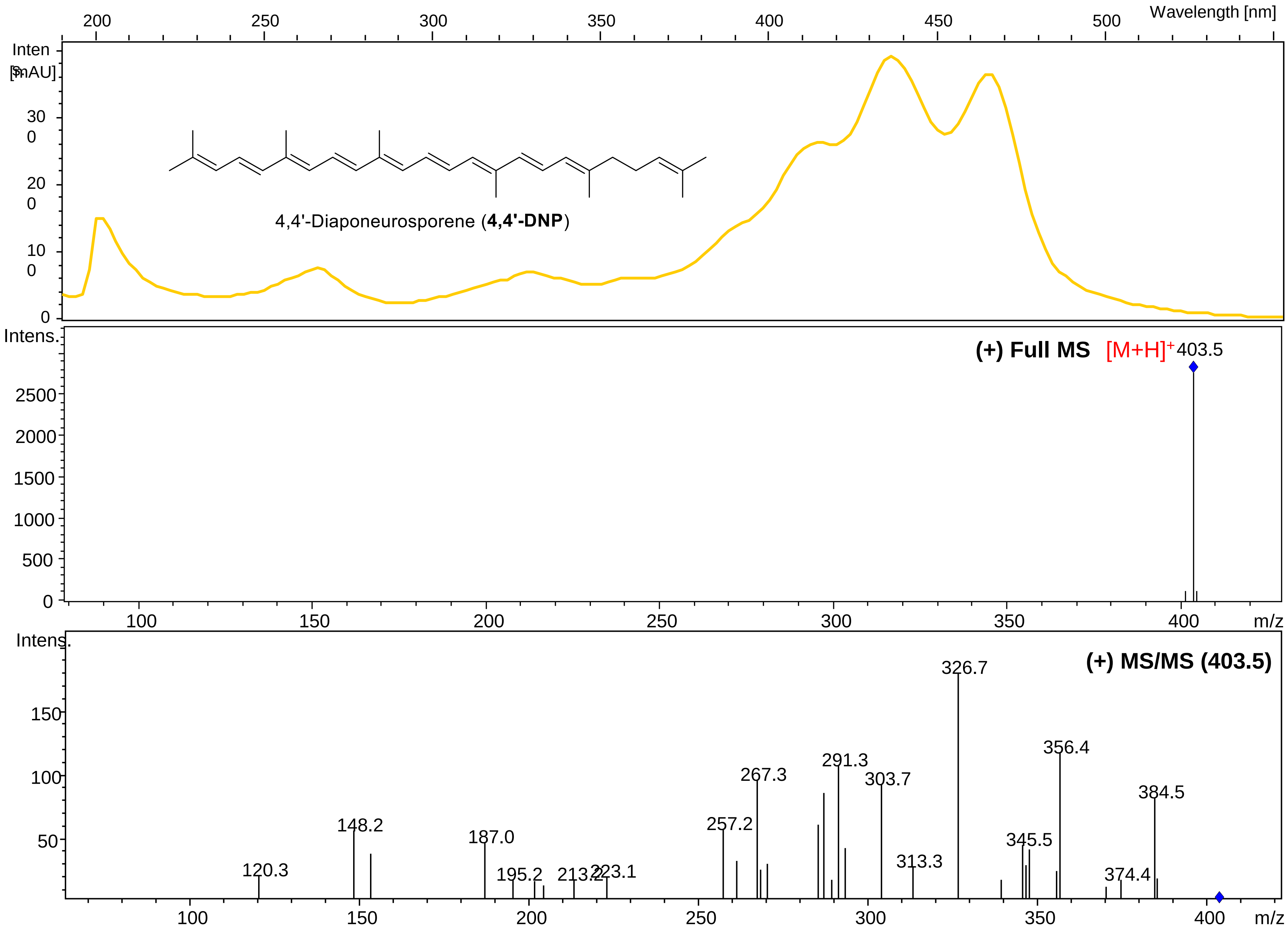


UV-visible, MS and MS/MS spectra for 4,4’-DNP in APCI (+).

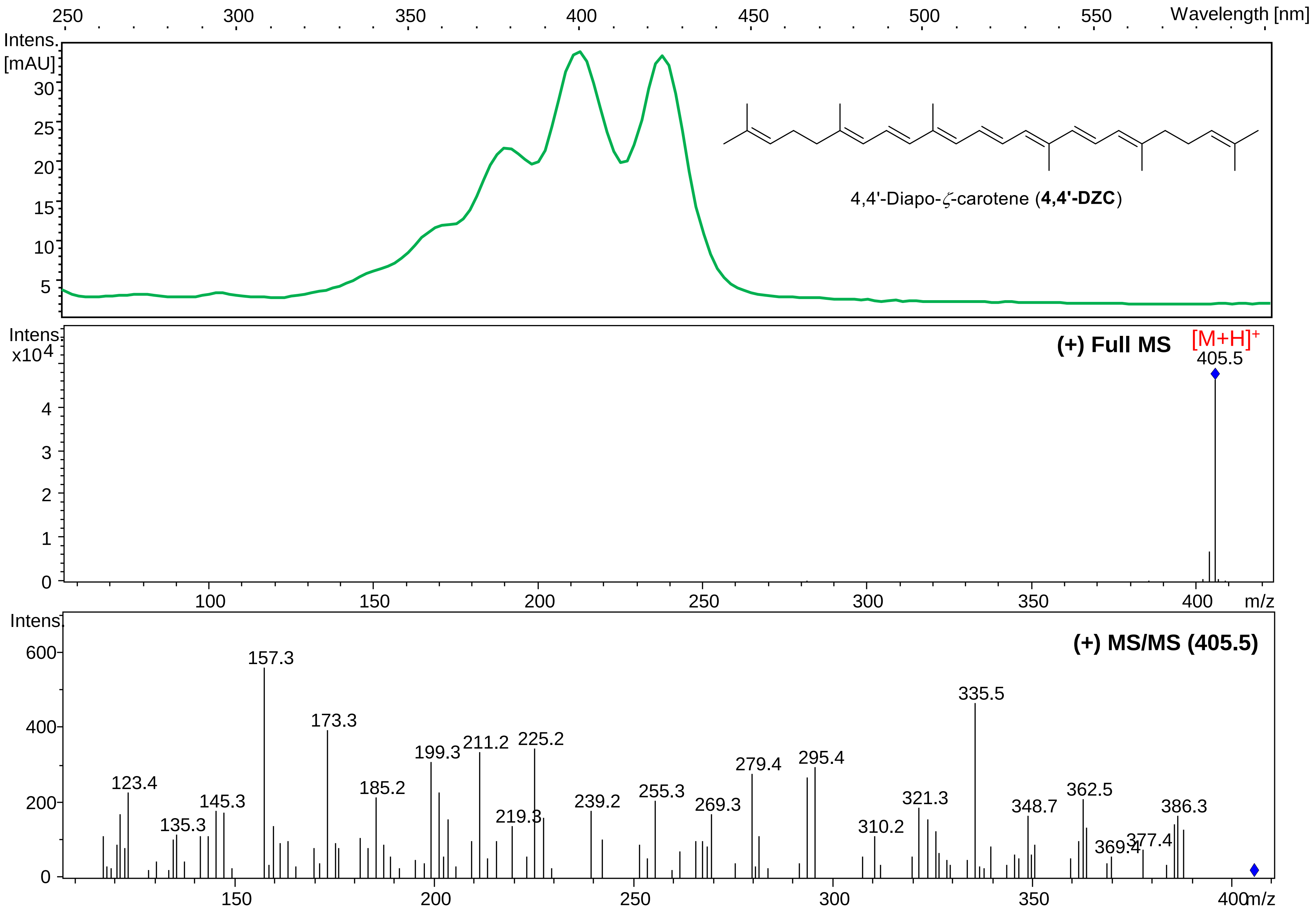


UV-visible, MS and MS/MS spectra for 4,4’-DZC in APCI (+).

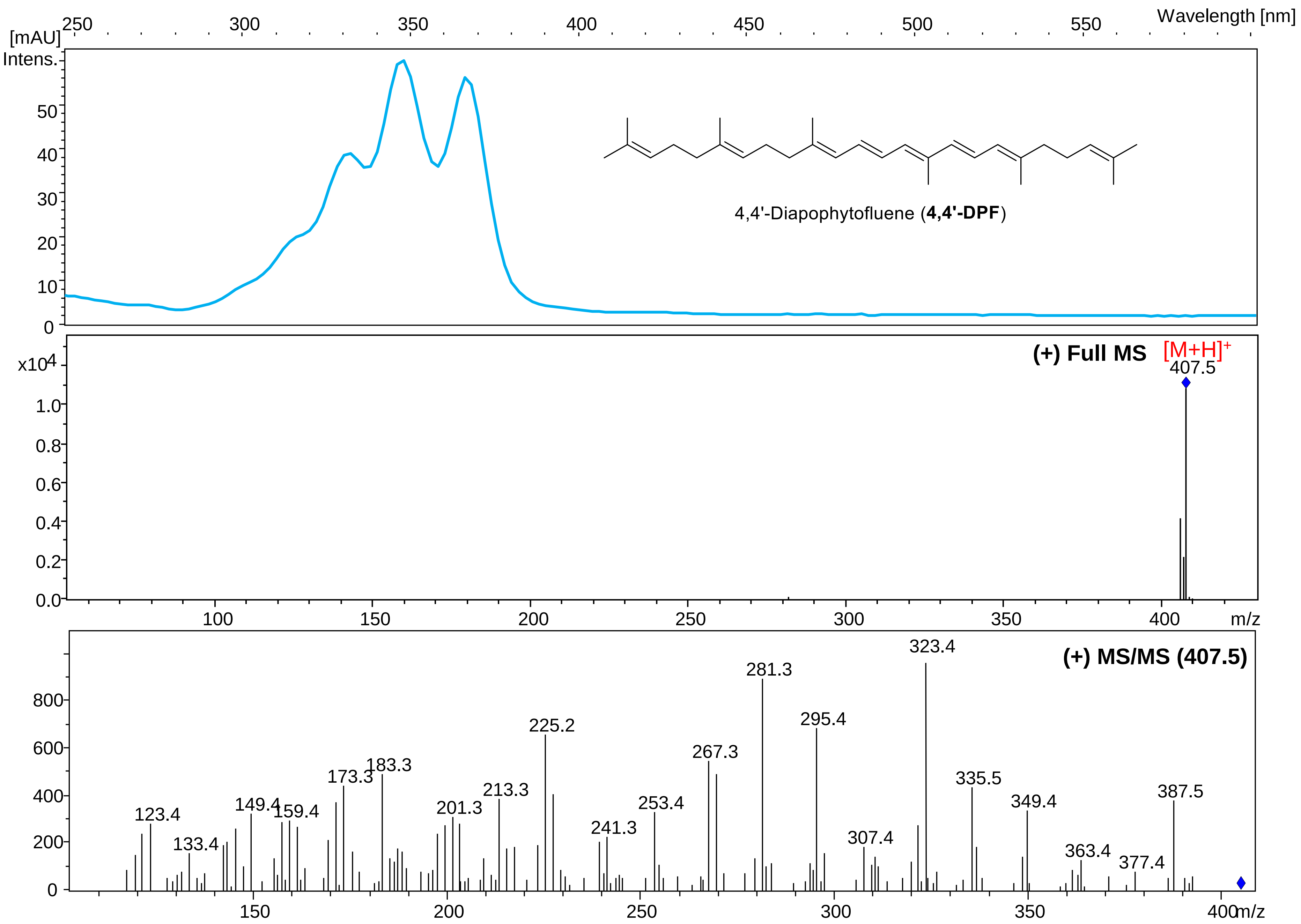


UV-visible, MS and MS/MS spectra for 4,4’-DPF in APCI (+).

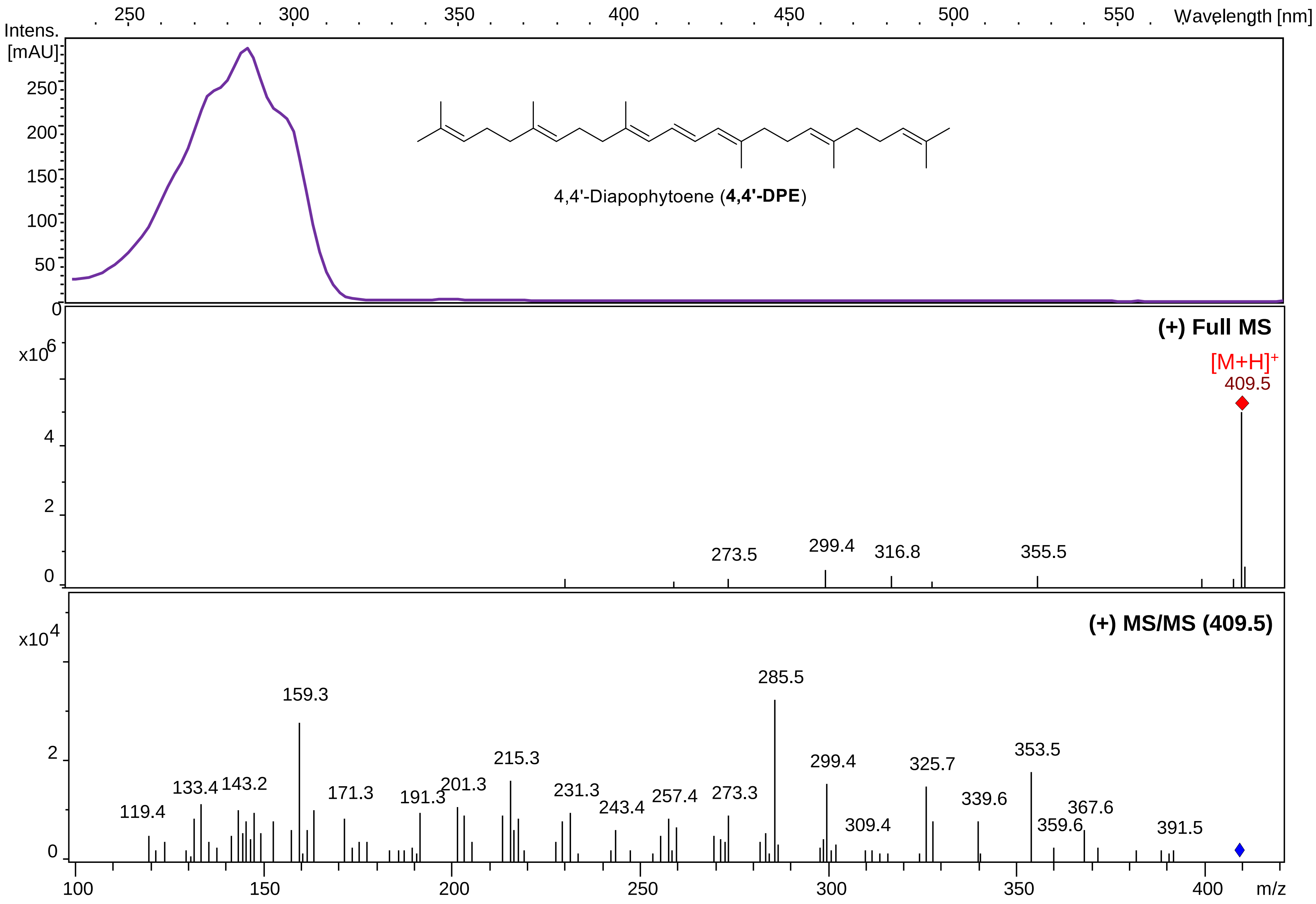


UV-visible, MS and MS/MS spectra for 4,4'-DPE in APCI (+).
