## Supplementary material for "Carotenogenesis of *Staphylococcus aureus*: new insights and impact on membrane biophysical properties": FigureS5

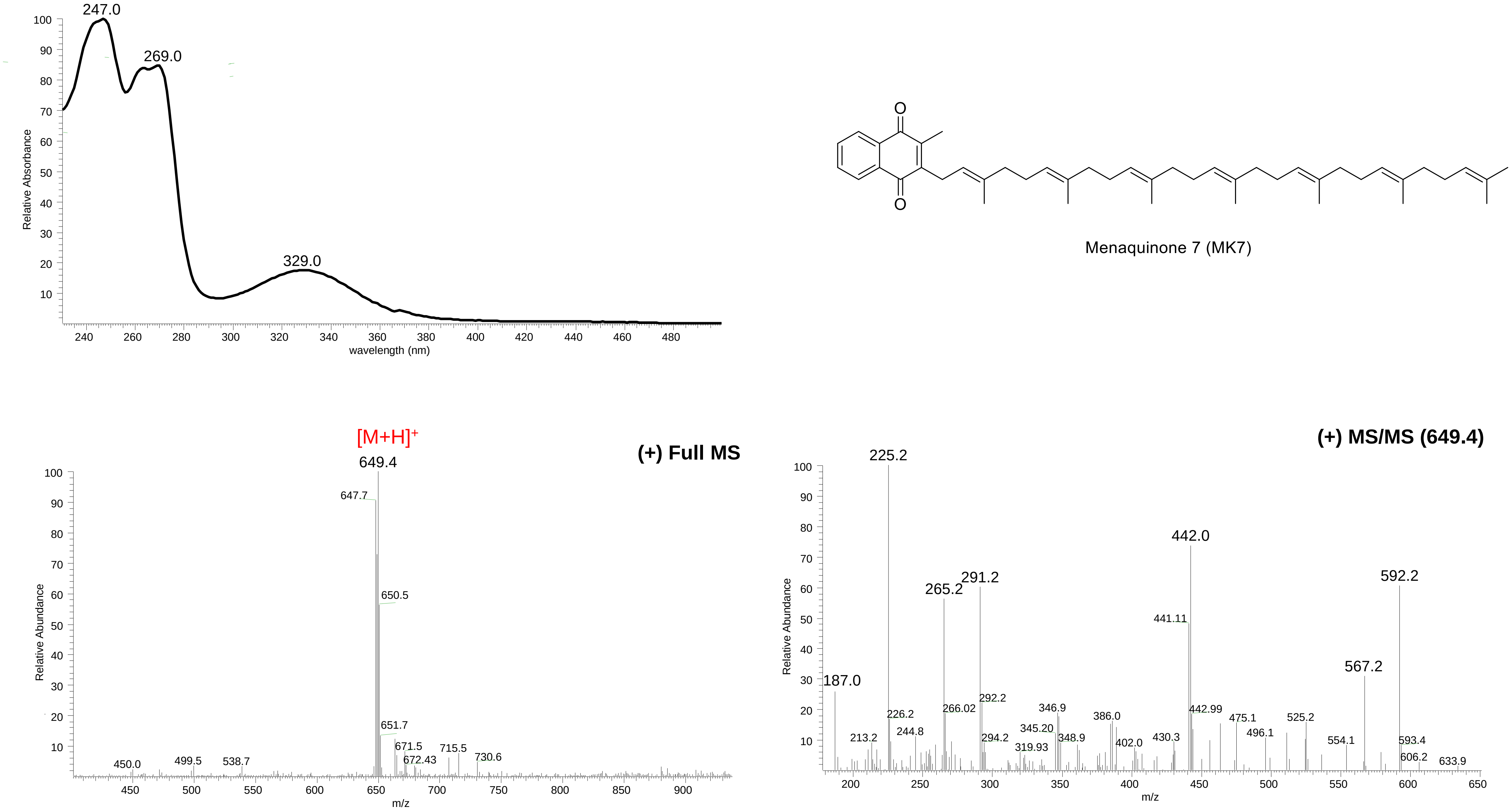


UV-visible, MS and MS/MS spectra for MK7 in ESI (+).

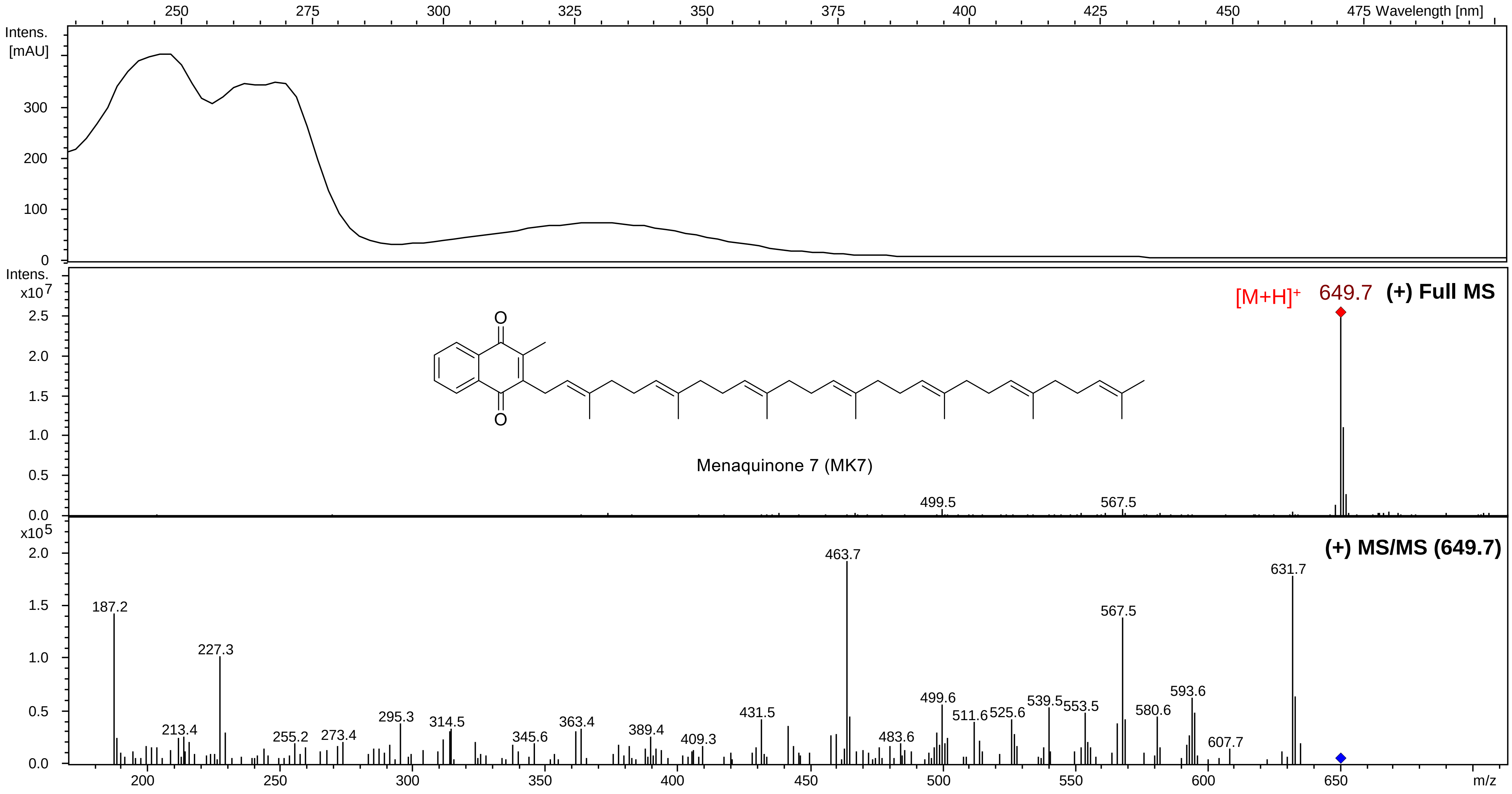


UV-visible, MS and MS/MS spectra for MK7 in APCI (+).

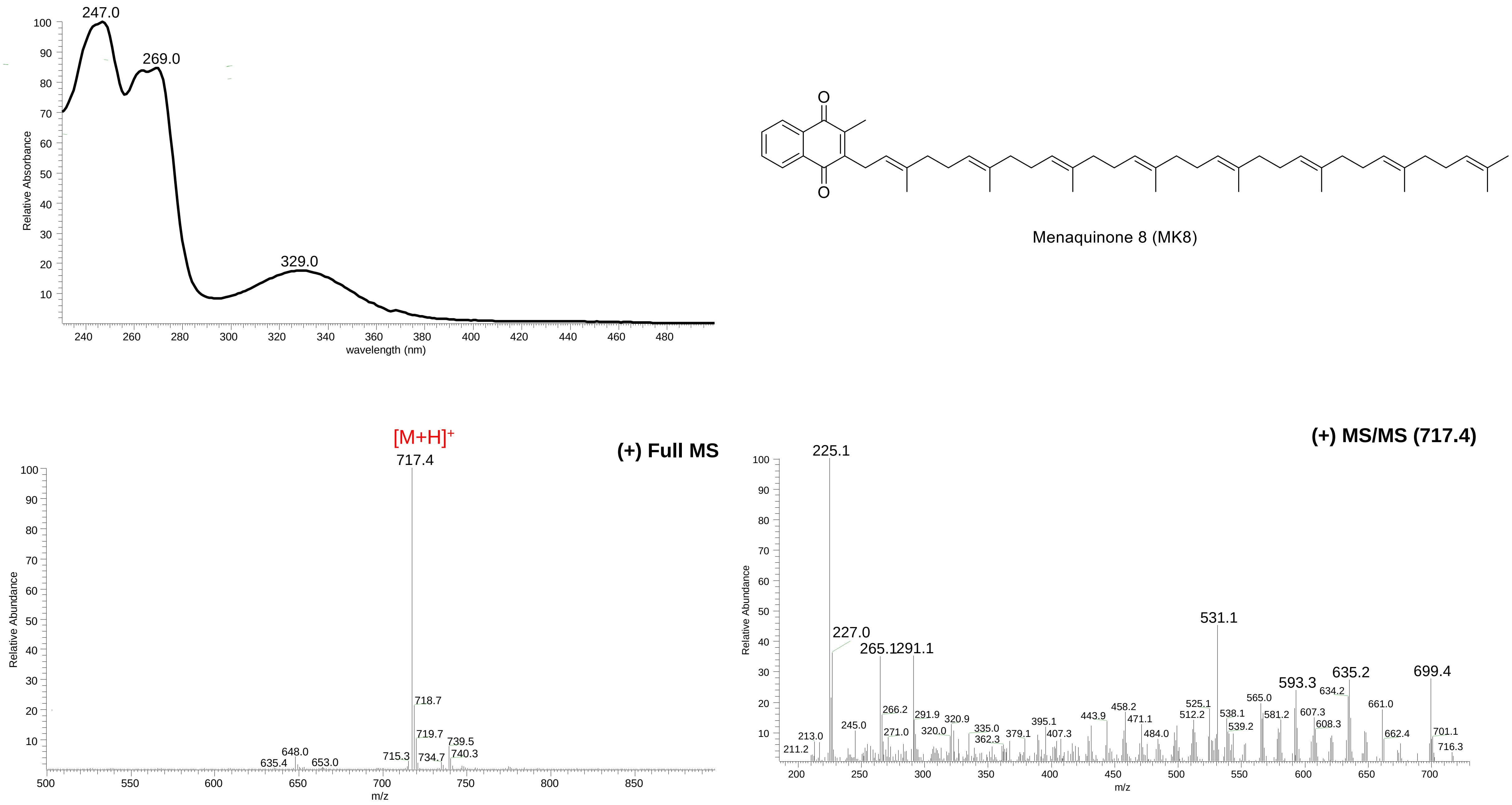

UV-visible, MS and MS/MS spectra for MK8 in ESI (+).

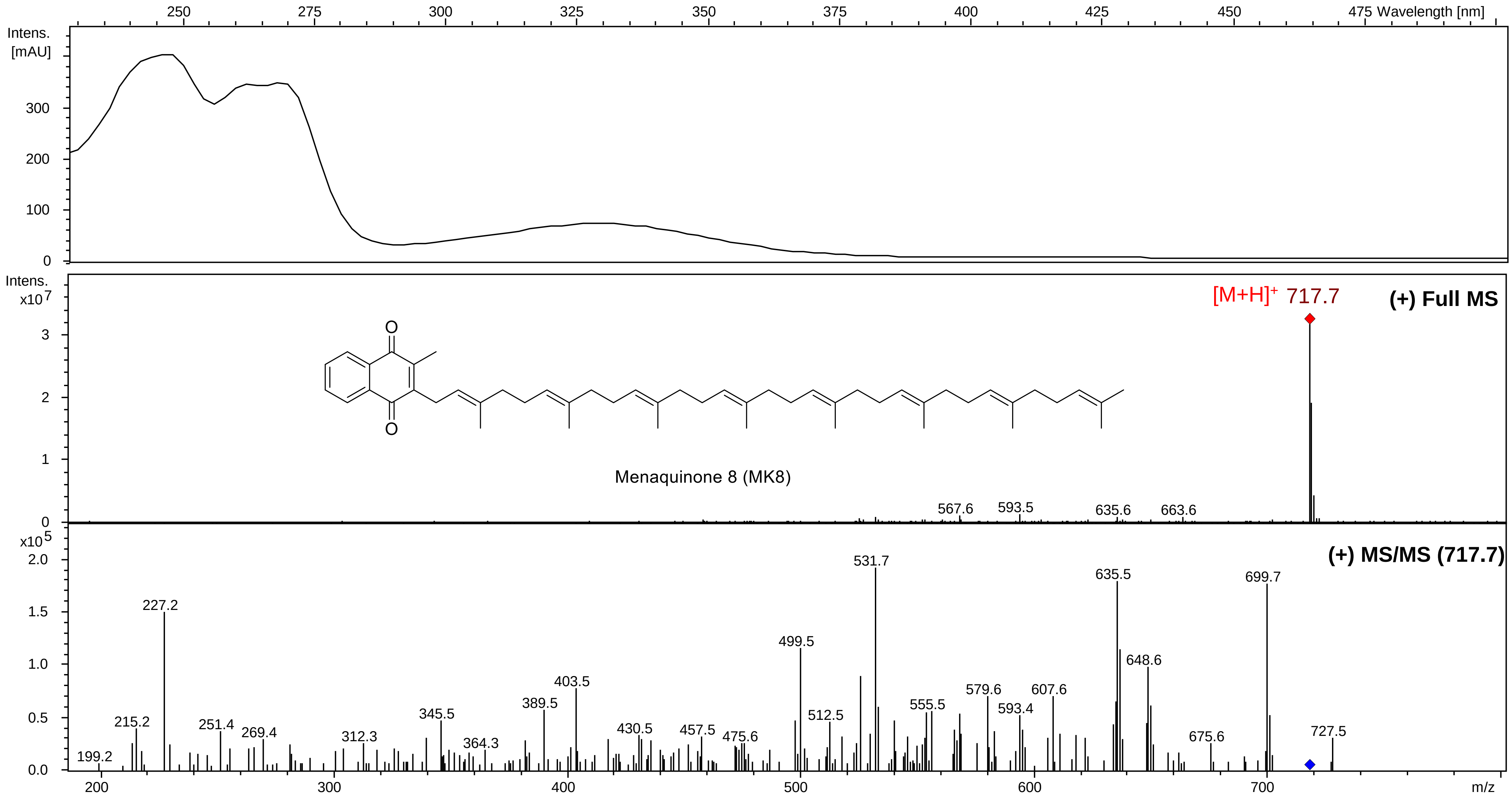


UV-visible, MS and MS/MS spectra for MK8 in APCI (+).

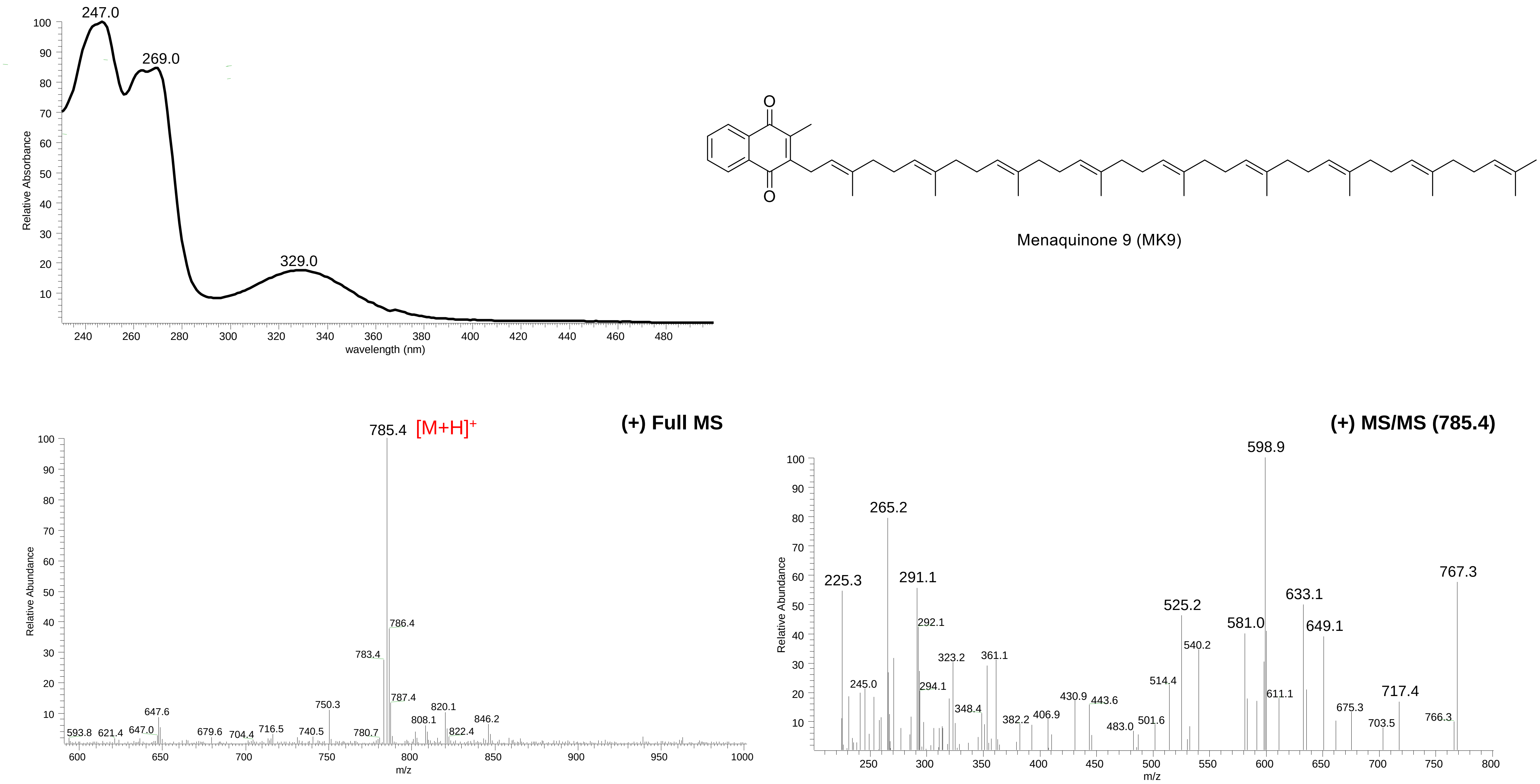


UV-visible, MS and MS/MS spectra for MK9 in ESI (+).

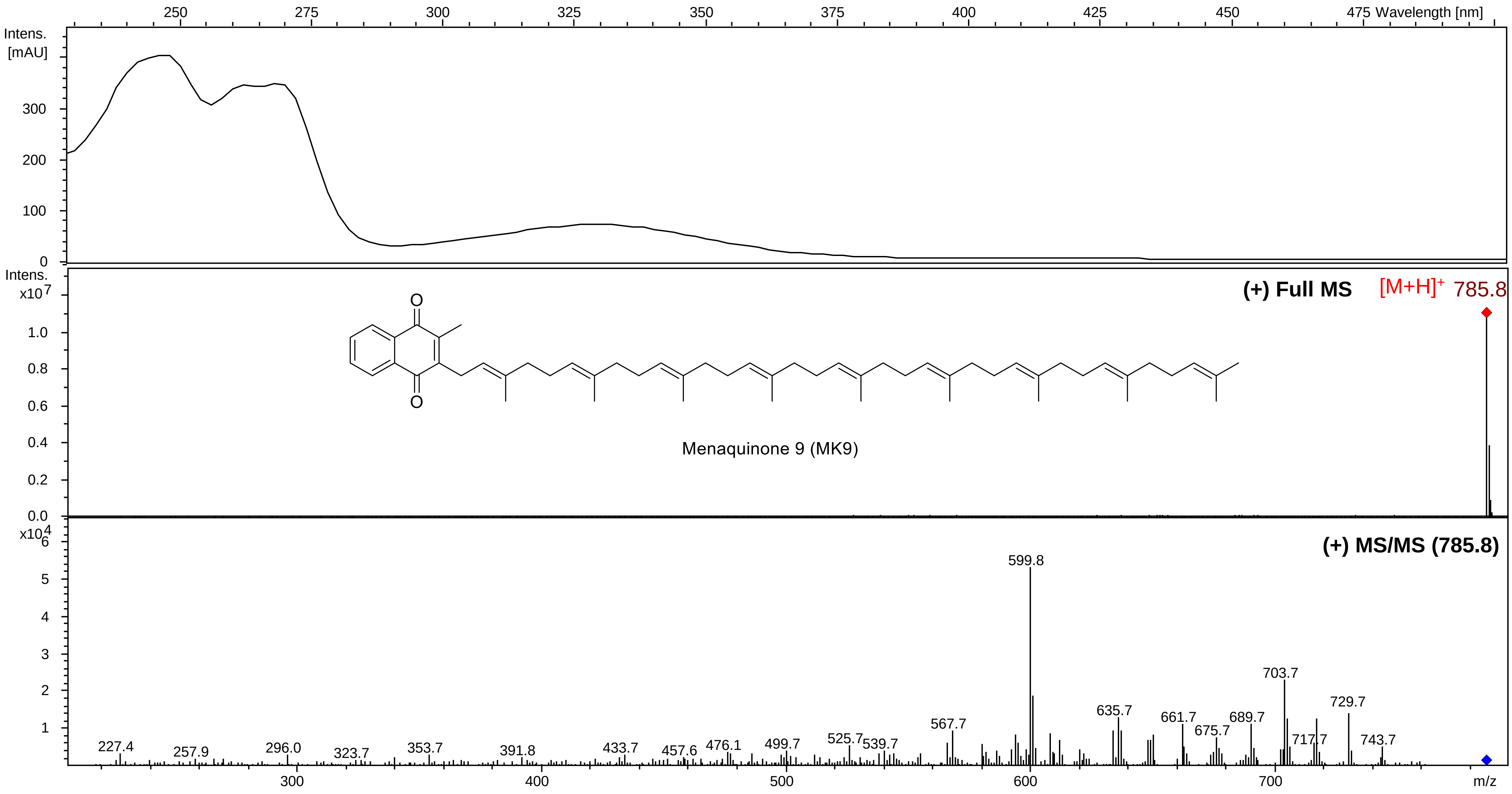


UV-visible, MS and MS/MS spectra for MK9 in APCI (+).
